# PKA disrupts adult ovarian identity circuits through direct inhibition of TRIM28-chromatin binding

**DOI:** 10.64898/2026.09.30.755598

**Authors:** Florian Chaleil, Lionel Le Gallic, Isabelle Sahut-Barnola, Joris Mordier, Stéphanie Le Gras, Sylvain Roumeau, Céline Keime, Cyril Djari, Antoine Vigier, Léna Badoil, Pierre Val, Jean-Christophe Pointud, Constantine A. Stratakis, Florence Cammas, Antoine Martinez, Francis Poulat, Anne Marie Lefrançois-Martinez

**Affiliations:** Institute of Genetics, Reproduction & Development (iGReD), CNRS, Inserm, University of Clermont-Auvergne, France; Service d’Endocrinologie, Centre Hospitalier Universitaire Gabriel Montpied, Clermont-Ferrand, France; Institute of Human Genetics (IGH), CNRS UMR9002, University of Montpellier; Department of Genetic Medicine and Development, University of Geneva, 1211 Geneva, Switzerland; Institute of Genetics and molecular and cellular biology, Illkirch-Graffenstaden; Center for Regenerative Therapies TU Dresden (CRTD), Technische Universität Dresden, Dresden, Germany; Hospital for Endocrine Surgery Research Center, HCA Florida Healthcare, HCA Healthcare Research Institute, Tampa, Florida, USA; Department of Pediatrics, University of Athens Medical School, Athens, Greece; Human Genetics & Precision Medicine, IMBB, FORTH, Heraklion, Greece

**Keywords:** Gonadotropic hormones, PKA, hypo-SUMOylation waves, granulosa cells, FOXL2, TRIM28 phosphorylation

## Abstract

Functional ovarian plasticity depends on the granulosa cell (GC) fate determination, differentiation, oocyte support and steroid production. After birth, GC identity must be maintained in differentiating follicles, while from puberty, ovarian plasticity is cyclically challenged by gonadotropic hormones. TRIM28-FOXL2 interactions form a key chromatin component maintaining ovarian identity programs and repressing testis-specific genes. This TRIM28 role depends on its E3-SUMO-ligase activity, suggesting SUMOylation in GC can be regulated by transient hormonal cues, although hormonal involvement in modulating GC identity remains to be established. Here, we demonstrated the *in vivo* interplay between gonadotropic-PKA-signaling and hypo-SUMOylation waves underlying ovarian GC plasticity. *In vitro* TRIM28 mutagenesis and phospho-proteomics demonstrated direct PKA inhibitory control on TRIM28. This prompted us to highlight SUMOylation alterations in Carney Complex ovarian lesions resulting from chronic PKA activity from rare genetic mutations. Using mutant models, transcriptomics and TRIM28/FOXL2-ChIP-Seq analyses, we specified the *in vivo* PKA inhibitory mechanisms on TRIM28/FOXL2 chromatin binding and its consequences on adult ovarian somatic cell identity. Together, our data support an integrative model where gonadotrophin pulses create transient windows of PKA-mediated TRIM28 phosphorylation, hypo-SUMOylation, and FOXL2–TRIM28 destabilization. This positions SUMOylation dynamics as central integrator of hormonal signaling and epigenetic identity maintenance in the ovary.

## INTRODUCTION

The mammalian ovary is a highly dynamic organ that undergoes important morphological changes during its development until sexual maturity, followed throughout the reproductive life by regular cyclical modifications in cellular differentiation processes and endocrine activities. Cyclic ovarian function produces mature female germ cells and sex hormones in follicles composed of an oocyte enclosed by granulosa cells (GC) and theca cells. Ovarian plasticity and activity depend on GC fate determination, proliferation, differentiation, and on their follicular role in oocyte support and steroid production, as GCs respond to paracrine and transient hormonal cues, growing and differentiating throughout folliculogenesis, ovulation and beyond. Genetic and genomic advances using mouse models established that WT1-KTS and FOXL2 factors together with WNT4 and RSPO1 signals form a core regulatory network ensuring GC specification and ovarian development while opposing DMRT1 and SOX9 autosomal factors that act with SRY for mammal testis differentiation^1^ (^2^for review). However, the finding that after birth, GC identity must be maintained by FOXL2 in conjunction with estrogen signaling in differentiating follicles provided the basis for molecular circuits required to ensure ovarian identity throughout reproductive life^3,4^, whilst from puberty, ovarian functional plasticity is cyclically challenged by FSH and LH. This concept has been confirmed by recent research demonstrating that TRIM28 interacts with FOXL2 and forms a key component of chromatin complexes maintaining ovarian identity programs and repressing testis-specific genes^5^. This latest study also showed that TRIM28 role in ovarian identity maintenance depends on its E3-SUMO ligase activity and demonstrated that epigenetic mechanisms involving SUMOylation dynamics interact with key sex regulators to ensure GC identity in adult ovaries, suggesting SUMOylation in GC is subjected to regulation throughout follicle development and maturation.

SUMOylation consists in the post-translational modification of target proteins by covalent addition of SUMO 1–3 peptides to specific lysine residues. It involves a sequential action of the E1 heterodimer SUMO-activating enzymes SAE1-2, the unique E2 enzyme UBE2I/UBC9 for SUMO conjugation to the lysine target through an isopeptide bond, and several E3 SUMO ligases including TRIM28 that increase efficiency of the conjugation step. The dynamics and reversibility of the reaction are ensured by SUMO-specific proteases of the SENP and DESI family that cleave and recycle SUMO peptides from their substrates (^6^for review). SUMOylation alters the interactions of target proteins with their partners, the latter often have one or more SUMO-interaction motifs (SIMs) that enhance recognition of SUMO-conjugated proteins^7^. The consequence of SUMOylation includes changes in cellular localization, protein activity, or protein stability. SUMOylation has emerged as a critical mechanism for modulating fundamental cellular and developmental processes whose dysregulations are implicated in severe diseases such as neurodegeneration, cancer or infection^8^. Further studies have highlighted the pivotal role of SUMOylation dynamics in key stages of early embryonic development^9^ and tissue morphogenesis^10^. The strict control of SUMOylation levels enables differentiation, cellular identity maintenance^11–13^ and homeostasis in several cell lineages^14,15^ including adrenocortical cells^16^ and adipocytes^17,18^, establishing SUMOylation as an emerging guardian of cell identity.

One of the major roles of SUMOylation cycle in the nucleus is the control of transcription processes through modification of trans-acting factors and chromatin-associated proteins^19–22^. In ovarian somatic cells, transcription factors (TF) such as LRH1/NR5A2, SF-1/NR5A1, and FOXL2 involved in development and function of steroidogenic tissues are regulated by SUMOylation^23–27^. Steroid nuclear receptors are also targeted by SUMOylation, including Estrogen receptors α and β^28–30^, which generally results in a decrease of their transcriptional activity^28^. A distinctive characteristic of SUMO regulation of transcriptional repression is that it is controlled in part by phosphorylation events caused by direct interference with SUMO moiety conjugation to target TFs^31^. Notably, progesterone receptor isoforms PR-A and PR-B are blocked by Progestins (R5020 and RU486)-induced SUMO-conjugation on Lys388 whereas Ser294 phosphorylation (through MAP3K or CDK2 action) selectively blocks PR-B SUMOylation, derepressing PR-B transcriptional activity^32^. Alternatively, phosphorylation events can indirectly repress SUMOylation by modulating expression or activity of SUMO machinery. Our work has previously demonstrated that SUMOylation of nuclear proteins is acutely and negatively regulated in the human and murine adrenal cortex by ACTH/PKA signaling pathway, through the simultaneous downregulation of SUMO-conjugating enzymes and upregulation of SUMO protease expression^33^. This ACTH/PKA-induced hypo-SUMOylation promotes adult adrenal cortex plasticity programs, allowing functional cortical zonation and stress responsiveness^16^.

In the ovary, PKA is a central integrator of FSH and LH gonadotropic signals which trigger the growth and maturation of the ovarian follicle as well as the ovulation process, alongside the functional reprogramming of GC into luteal cells^34–39^. In this context, the physiological relevance of direct or indirect PKA connections to several follicular nuclear factors as well as to histones and chromatin modulators has been firmly demonstrated^35,40–44^. However, PKA involvement in modulating molecular circuits that maintain postnatal GC identity remains to be established.

In this work, we investigated the interplay between pituitary gonadotropin hormones (PGH) and SUMOylation dynamics in ovarian functional plasticity by exploring *in vivo* and *ex vivo* the mechanisms by which PGH/PKA signaling interacts with SUMOylation machinery in GC of wild-type mice ovaries. We next developed *in vitro* PKA-GOF (gain-of-function) systems, TRIM28 directed mutagenesis and TRIM28 phospho-proteomic analyses to specify biochemical and functional interactions between PKA and E3-SUMO-ligase TRIM28. These data prompted us to investigate alterations in SUMOylation in rare ovarian tissue lesions from Carney Complex patients who exhibit genetically-induced chronic PKA activation. Finally, we utilized transcriptomic analyses conducted on various mouse models – including those exhibiting PKA-GOF in gonadal somatic cells –to define the *in vivo* links between PKA activity and the TRIM28/FOXL2 nuclear complexes, as well as their implications for the maintenance of adult ovarian somatic cell identity. We then performed Proximity Ligation Assay, salt-induced chromatin dissociation and Chromatin Immuno-Precipitation Sequencing (ChIP-Seq) to determine the pivotal role of PKA-TRIM28 interactions that influence sex-maintaining gene landscapes of ovarian somatic cells.

## RESULTS

### Gonadotropic hormones induce transient ovarian hypo-SUMOylation waves *in vivo* and modulate SUMO-player gene transcription *via* PKA signaling

To determine whether the gonadotropin-PKA coupling modulates the dynamics of SUMOylation in the ovary, we first examined the impact of PGH on SUMOylation patterns *in vivo*. The hypothalamic-pituitary-gonadal (HPG) axis activity in 25-day-old wild-type (WT) mice was synchronized using the GnRH antagonist cetrorelix (CTX), then driven using gradual exposure to equine chorionic gonadotrophin (eCG) followed by human chorionic gonadotrophin (hCG), which activate the murine FSH and LH receptors, to ensure the follicle maturation and the ovulatory signals, respectively^45^ (Fig. 1a).

**Figure 1:**
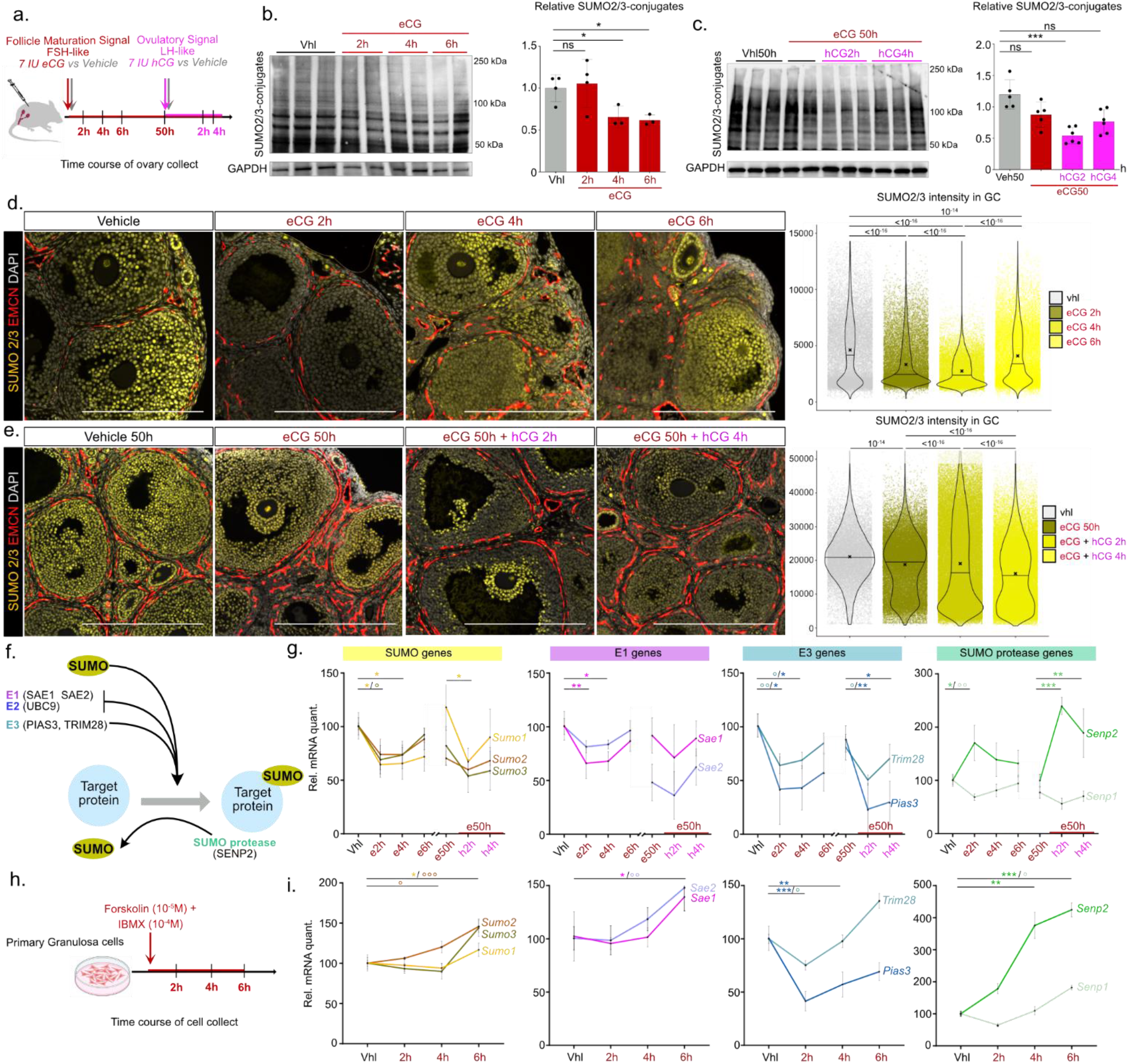
Gonadotropic hormone-induced-PKA activity exerts transient hypo-SUMOylation waves *in vivo*, in ovarian follicles and modulates transcription of SUMO-player genes in ovarian somatic cells. **a.** Schematic experimental design of gonadotrophin stimulation. Following HPG synchronization with the GnRH inhibitor CTX, OF1 WT mice were divided into groups. The first group received eCG, and ovaries were collected 2h, 4h, 6h, or 50h later. The second group received hCG 50h after the first injection of eCG, and ovaries were collected 2h or 4h later. Control ovaries received vehicle (vhl) and were collected at 2h or 50h. **b. c.** Time course effects of gonadotrophins on the accumulation of ovarian SUMO2/3-conjugated-proteins analysed by WB with SUMO2/3-antibody and SUMO2/3 signal quantification. GAPDH signal was used to normalize the quantification of SUMO signals. Statistical analysis was performed using two-sited Mann-Whitney U test. **d. e.** Co-immunofluorescence labelling of SUMO2/3 (yellow) and endomucin (endothelial cells) (red) in WT treated ovaries. The quantification graphs represent the quantification of SUMO2/3 intensity in GC nuclei of ovaries from 3 or 4 different mice for each hormonal treatment. *P* value was determined with a Kruskal–Wallis test with multiple comparisons using Dunn’s test. Scale bar: 200µm. **f.** Schematic representation of the SUMOylation process. SUMO1, SUMO2, and SUMO3 peptides are conjugated to target proteins by E1, E2, and E3 enzymes and removed by proteases such as SENP2 or DESI1. **g.** RT-qPCR analyses of SUMOylation pathway gene expression. Quantification histograms represent means ± SD. n= 5/6 ovaries *per* genotype. Statistical analyses were conducted by a Kruskal–Wallis test with multiple comparisons using Dunn’s test. \**P*<0.05, \*\**P*<0.01. **h.** Experimental design of primary GC treatments before RNA analyses. Murine primary GC were treated with forskolin (FK) (10^-5^ M) and 3-isobutyl-1-méthylxanthine (IBMX) (10^-5^ M) for 2h, 4h or 6h or with vehicle (Vhl) (DMSO) for the longest treatment time. **i.** RT-qPCR analyses of SUMOylation player gene expressions in primary GC. Quantification histograms represent means ± SD. n= 5/6 ovaries *per* genotype. Statistical analyses were conducted by a Kruskal– Wallis test with multiple comparisons using Dunn’s test. \**P*<0.05, \*\**P*<0.01, \*\*\**P*<0.005.

Western blot analyses of SUMO2-3-conjugates accumulation in whole ovaries revealed that significant global hypo-SUMOylation of ovarian proteins occurred as early as 4 hours after eCG exposure (Fig. 1b) and which reappeared transiently for less than 2 hours, 2 hours after exposure to hCG, in ovaries that had previously been stimulated for 50 hours with eCG (Fig. 1c). In contrast, we did not observe any change in SUMO1-conjugates levels in response to gonadotropin treatments (Supplementary Fig. S1a-b).

We next performed immunohistochemical analyses on tissue sections to identify the ovarian compartments where PGH-induced hypo-SUMOylation was occurring. SUMO2/3 immunolabeling was widely distributed in most GC nuclei in the follicular compartment and in stromal cells in CTX-synchronized ovaries. In contrast, the staining began to fade noticeably and rapidly in the synchronized ovaries exposed to eCG for 2 and 4 hours. From 6 hours of exposure, SUMO2/3 nuclear staining was gradually restored in follicular and in stromal cells (Fig. 1d). Similar changes were observed in response to hCG treatment kinetics (Fig. 1e). Quantification of labeling intensity adjusted to GC across all follicles confirmed significant waves of variation in SUMO2/3-conjugate levels in GC nuclei in response to PGH.

As SUMOylation dynamics result from a cascade of enzymatic reactions (Fig. 1f), we analyzed the expression levels of genes encoding SUMO players in whole ovaries in response to gonadotropic stimulations (Fig. 1g). RT-qPCR analyses revealed that gonadotrophin stimulation with eCG, which governs follicular maturation, exerted a rapid (as early as 2 hours) and transient inhibitory control over transcript levels of *Sumo1*, *Sumo3*, *Sae1*, *Pias3* and *Trim28* genes, as well as a rapid activating effect on *Senp2* expression. The ovulatory signal induced by hCG treatment exerted similar rapid effects on transcript levels of *Sumo1*, *Pias3*, *Trim28*, and *Senp2*, indicating that transient hypo-SUMOylation of ovaries in response to gonadotropins is broadly concomitant with inhibition of pro-SUMOylating gene expression and activation of de-SUMOylating gene expression.

Under normal physiological conditions, cAMP-PKA signaling is one of the major mediators of FSH and LH controls in the ovary. To focus on PKA role in SUMOylation dynamics, we analyzed gene expression in primary GC cultures exposed to forskolin (FK) and isobutyl methyl xanthine (IBMX) to stimulate and stabilize cAMP/PKA signaling (Fig. 1h). RT-qPCR revealed that only *Trim28* and *Pias3* transcripts fell rapidly and transiently within the first 2 hours of treatment, while *Senp2* rose rapidly throughout the treatment period, in contrast to other SUMO protease transcripts, which increased more slowly (Fig. 1i, Supplementary Fig. S1c).

Altogether, these data indicate that in GC, cAMP/PKA signaling exerts rapid coordinated transcriptional control that inhibits *Trim28* and *Pias3* while activating *Senp2*. However, these transcriptional kinetics alone cannot account for the simultaneous variations in SUMO-conjugated protein accumulation, suggesting additional post-transcriptional PKA controls regulate SUMOylation dynamics.

### PKA directly phosphorylates TRIM28^S473/S489^ to inhibit its E3-SUMO ligase activity

TRIM28 is indirectly connected to the genome through its N-terminal domain interacting with zinc finger TFs^46^. In its C-terminal domain, it contains a plant homeodomain (PHD) and a bromodomain (Br) (Fig. 2a) involved in recognizing histone modifications^47^. Notably, TRIM28 possesses E3-SUMO ligase activity, preferentially using SUMO2-paralogs as substrates^48^. Its activity is mediated by its PHD domain, which functions similarly to SUMO ligase domains of PIAS proteins^49–51^. When SUMO-conjugated by UBC9, TRIM28 acts as an intramolecular E3-SUMO ligase on 6 lysines in its PHD and bromo domains, then recruits the NuRD complex and SETDB1 methyltransferase, enabling removal of histone acetylation marks and addition of repressive H3K9me3 marks. The binding of heterochromatin protein 1 (HP1) helps stabilize the complex by binding to H3K9me3 marks, inducing heterochromatin environment formation^52,53^. TRIM28 interaction with HP1 is compromised by kinase-induced-phosphorylation on serine residues, notably Ser473, located close to the PHD domain^52,53^. Given that *Trim28* inactivation or PHD domain deletion in follicles at puberty disrupts GC identity maintenance^5^, we hypothesized that TRIM28-E3-SUMO ligase activity might be modulated by direct PKA phosphorylation. To investigate whether PKA alters TRIM28 phosphorylation state, endogenous TRIM28 was immunoprecipitated from HEK293T cells under basal conditions, after FK treatment, or following expression of constitutively active PKA catalytic subunit (PKA-Cα), and analyzed by phospho-proteomics (Fig. 2b). To maximize sequence coverage, parallel digestions with Lys-C and chymotrypsin were performed. These complementary approaches enabled mapping of the majority of TRIM28 phosphorylation sites. Within this mapped set, Ser473 and Ser489 were the two sites showing clear stimulus-dependent phosphorylation increases (Fig. 2c).

**Figure 2:**
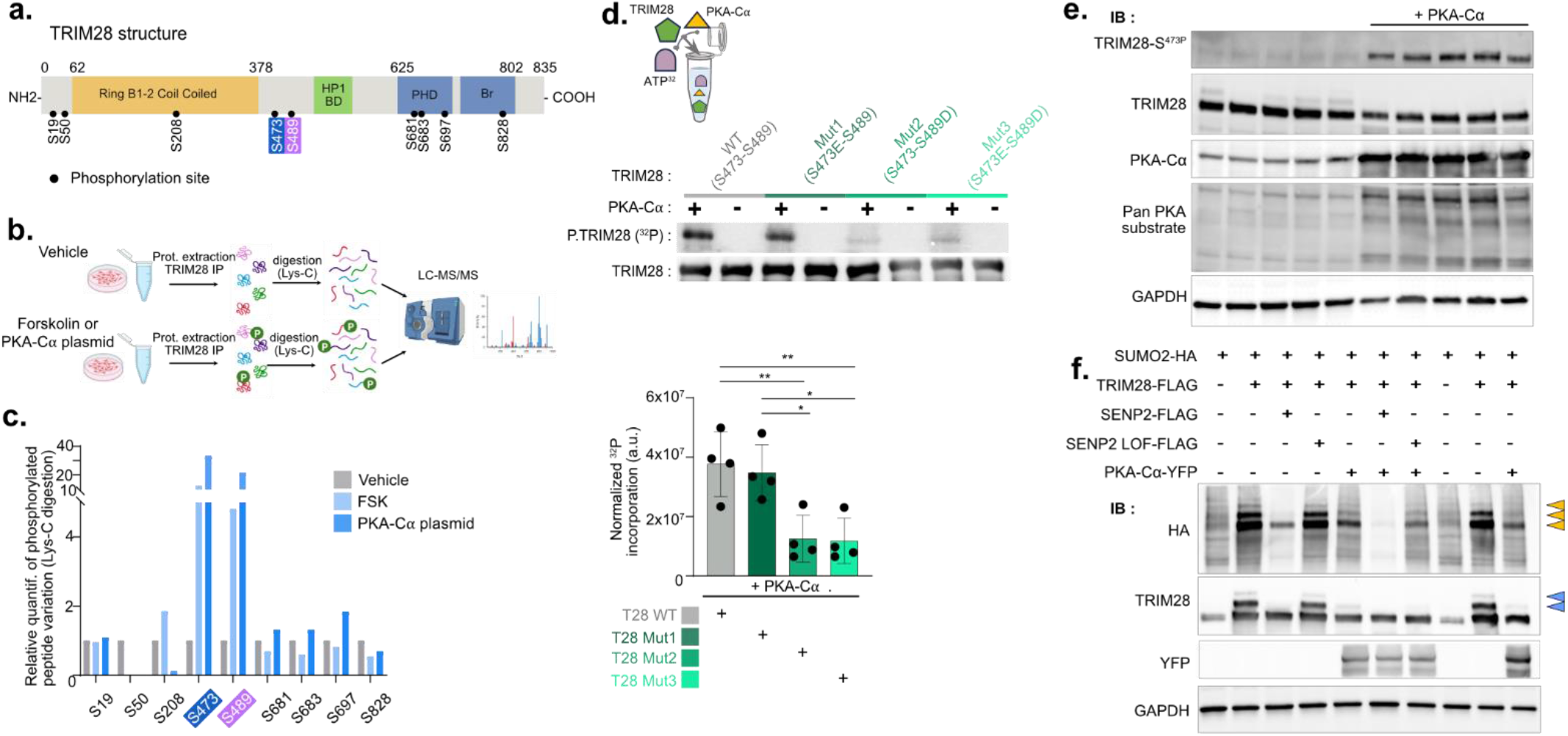
PKA exerts a direct inhibitory control of the TRIM28-E3SUMO-ligase activity by phosphorylating the TRIM28-S^473^ and -S^489^. **a.** Schematic of the TRIM28 with three main domains: RBCC, HP1 binding and PHD SUMO domains. The serine residues identified by mass spectrometry were listed as phosphorylation sites. **b.** Phosphoproteomics workflow: HEK293T cells were subjected to three treatments: untreated control or PKA stimulation (treatment of 50 µM forskolin (FSK) for 30 min, or transient transfection with a pcDNA-PKA-Cα expression vector for 24h). After protein extraction and endogenous TRIM28 immunoprecipitation, bead-bound proteins were digested on-bead using Lys-C to maximize TRIM28 sequence coverage and phosphosite detection. The resulting peptides were analyzed by nano LC-MS/MS. Peptides and phosphorylation sites were identified and relative phosphorylation levels were determined by comparing the chromatographic peak areas of phosphorylated and non-phosphorylated peptide species. **c.** Phosphoproteomic analysis of purified TRIM28. Purified TRIM28 was digested with Lys-C and analysed by LC-MS/MS. Phospho-site assignment was validated using AScore and -10logP confidence scores. Fraction was calculated as phosphorylated peptide intensity/(phosphorylated + non-phosphorylated peptide intensity). For each condition, a relative quantification of the variation in phosphorylated peptides compared to the vehicle condition was calculated. **d.** PKA sequentially phosphorylates TRIM28 at Ser473 and Ser489. Recombinant His-tagged wild-type (WT) TRIM28 and the phosphomimetic mutants S473E, S489D and S473E/S489D were incubated with the catalytic subunit of PKA in the presence of γ-32P-ATP. Upper panel: Representative phosphorimages (P-TRIM28) and T7 immunoblots (WB TRIM28). Lower panel: Quantification normalized of ^32^P incorporation (a.u., arbitrary units). Data are presented as mean ± SEM from n = 4 independent kinase assays. Statistical significance was determined by one-way ANOVA followed by Tukey’s multiple-comparison test. \**P*<0.05, ** *P*<0.01. **e.** WB analysis of TRIM28 phosphorylation in HEK293T cells transfected with or without expression plasmid encoding the PKA active catalytic unit Cα (PKA-Cα). Pan PKA substrate is potent and selective substrate peptide of PKA that is used to detect PKA activity. **f.** WB analysis of TRIM28 SUMOylation in HEK293T cells co-transfected with SENP2-FLAG, SENP2 LOF-FLAG and PKA-Cα-YFP. WB were revealed with HA, TRIM28, PKA-Cα-YFP, GAPDH antibodies.

Although Ser473 was a previously established TRIM28 phospho-site, its modification by PKA activation was not experimentally demonstrated. In our dataset, Ser473 was confidently quantified in Lys-C digests, where its occupancy increased 12.33-fold after FK stimulation and 30-fold following PKA-Cα overexpression. By contrast, this site was not recovered by chymotrypsin, most likely due to peptide-specific constraints (Supplementary Fig. S2c). These results indicated that Ser473 is responsive to cAMP/PKA pathway activation.

Ser489 represents a previously characterized phospho-site in TRIM28^54^, located within the HP1-binding region spanning amino acids 483–497 and overlapping the HP1 box (⁴⁸⁶PXVXL⁴⁹⁰)^52^. In chymotrypsin digests, Ser489 displayed a 4.9-fold occupancy increase after FK stimulation and 8-fold after PKA-Cα overexpression. The same trend was independently observed in Lys-C digests (4.6-fold and 20.37-fold, respectively). Finally, our analysis was consistent with NetPhos predictions pointing to these two serines among the most likely PKA targets (Supplementary Fig. S2d-e). Together, these data identify Ser489 as a novel, highly responsive PKA phosphorylation site and suggest that cAMP/PKA signaling may regulate TRIM28 function through a site positioned in the HP1-binding region.

To determine whether TRIM28 is directly phosphorylated by PKA, an *in vitro* kinase assay was performed using bacterially expressed and purified WT TRIM28 together with S473E, S489D, and S473E/S489D phosphomimetic mutants, purified PKA catalytic subunit, and γ-³²P-ATP.

WT TRIM28 was efficiently phosphorylated by PKA (Fig. 2d). The S473E mutant displayed WT-comparable ³²P incorporation, indicating the conformational change does not impair phosphorylation at a second residue, likely Ser489, and may even facilitate it. In contrast, S489D exhibited marked reduction in ³²P incorporation, comparable to the double mutant, which retained only residual phosphorylation. These results suggest phosphorylation of Ser473 promotes subsequent Ser489 phosphorylation, whereas Ser489 modification prevents Ser473 phosphorylation. Together, our data support a sequential mechanism where PKA phosphorylates TRIM28 in a defined order: first at Ser473, then at Ser489.

We next investigated endogenous TRIM28 phosphorylation using the available anti-TRIM28-S^473P^ antibody in H293 cells transfected with PKA-Cα-YFP. Western blotting revealed very low endogenous TRIM28-S^473P^ in empty vector controls, whereas higher levels were detected upon PKA-Cα-YFP overexpression (Fig. 2e). To specify PKA consequences on TRIM28-E3-SUMO ligase activity, we performed co-transfection experiments in HEK293T cells with Trim28-Flag, PKA-Cα-YFP, Sumo2-HA, and Senp2-Flag expression vectors. HA-immunodetection showed SUMO2-conjugates of TRIM28 as three bands of decreasing intensity at approximately 130, 160, and 190 kDa (Fig. 2f, yellow arrowheads) that were eliminated when WT-Senp2 was ectopically co-expressed. TRIM28-immunostaining confirmed the 130 and 160 kDa bands as main conjugates (Fig. 2f, blue arrowheads). Accumulation of all TRIM28-SUMO2-HA conjugates was reduced upon PKA-Cα-YFP overexpression, suggesting PKA-dependent repression of TRIM28 auto-SUMOylation. Altogether, these experiments establish that Ser473 and Ser489 are the main direct PKA targets.

### PGH induce transient TRIM28-S^473P^ accumulation and acute negative control of SUMOylation dynamics in ovarian GC

Despite lacking a TRIM28-S^489P^ antibody, and because our in vitro results suggest a concomitant phosphorylation by PKA on both residue we used anti-TRIM28-S^473P^ to investigate *in vivo* changes of TRIM28-S^473P^ accumulation in response to PGH treatments (Fig. 1a). WB analyses revealed a significant, relative, and transient increase in TRIM28-S^473P^ accumulation in whole ovary protein extracts as early as 2 hours after eCG and 2 hours after hCG treatment (Fig. 3a-b). Immunohistochemistry on ovary sections specified that these variations in TRIM28-S^473P^ immunolabeling mainly occurred in GC nuclei (Fig. 3c-d) and mirrored the disappearance of SUMOylation signal in the same time course (Fig. 1d-e).

**Figure 3:**
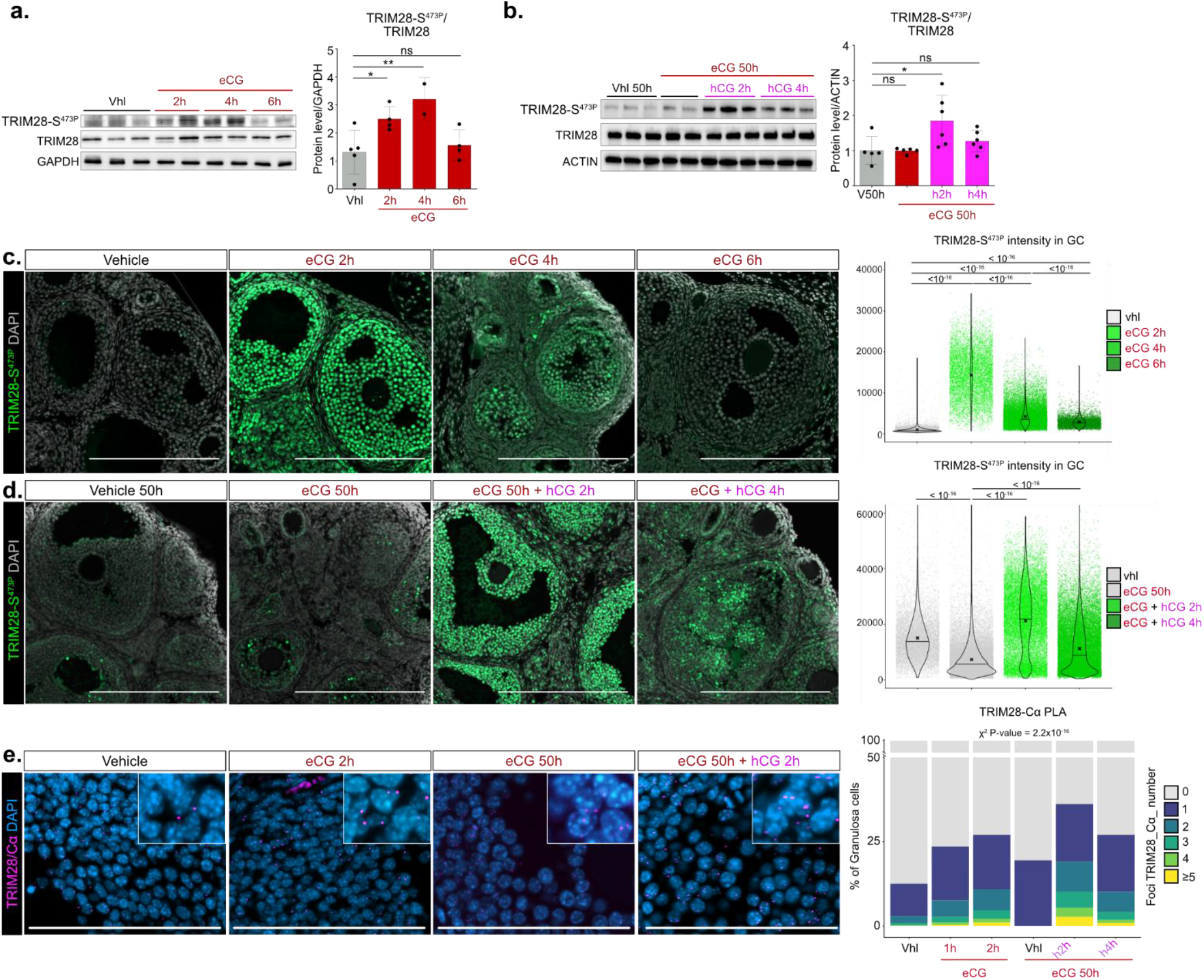
PGH exert an acute and transient negative control on ovarian granulosa cell SUMOylation dynamics *in vivo*, associated to PKA induced inactive TRIM28-S^473P^ increase of accumulation. **a. b.** Time course effects of gonadotrophins on the ovarian TRIM28-S^473P^ accumulation analysed by Western blot and quantification of the TRIM28-S^473P^/total-TRIM28 signal ratio. GAPDH or ACTIN signal was used to normalize the quantification of TRIM28-S^473P^ and TRIM28 signals. Statistical analysis was performed using two-sited Mann-Whitney U test. **c. d.** Immunofluorescence of TRIM28-S^473P^ in gonadotrophin treated ovaries. The graphs represent the quantification of TRIM28-S^473P^ immunostaining signal intensity in GC nuclei from ovaries from 3/4 mice *per* treatment. *P* value was determined with a Kruskal–Wallis test with multiple comparisons using Dunn’s test. Scale bar: 200µm. **e.** Proximity ligation assay (PLA) analyses of PKA interactions with TRIM28 in GC nuclei of gonadotrophin treated ovaries. Histograms represent proportion of GC containing the specified number of dots of interaction *per* nucleus of treated ovaries. To assess whether the distribution of cells differs significantly between the groups a χ² test of independence was assessed. n=3 ovaries *per* treatment. Representative histological images of all conditions are shown in Supplementary Figure S3. \**P*<0.05, \*\**P*<0.01, \*\*\**P*<0.005.

Since our *in vitro* analyses highlighted direct physical and functional interaction between active PKA-Cα and TRIM28, we visualized their interaction in GC using *in situ* proximity ligation assay (PLA), which enables identification of both stable and transient protein/peptide interactions. We confirmed a significant increase in GC numbers exhibiting PLA signals and in the number of interaction foci between PKA-Cα and TRIM28 *per* GC nucleus in response to eCG as early as 1 hour, peaking at 2 hours, and again at 2 hours after hCG (Fig. 3e, Supplementary Fig. S3a). No signal was detected 50 hours after eCG, indicating that PKA-Cα-TRIM28 physical interaction is transiently enhanced in response to PGH treatment.

Before puberty, HPG axis function results in transient PGH production episodes, beginning at the end of the fetal period and recurring in infantile mice with isolated acute FSH increases, culminating at PND14^55^. This mini-puberty period gives rise to a transient ovarian response^56,57^. We took advantage of this physiological context to examine spontaneous gonadotrophin variation effects on SUMOylation dynamics and associated GC TRIM28-S^473P^ accumulation during postnatal ovarian ontogenesis. Our IHC analyses revealed strong SUMO2/3-conjugate immunostaining in GC nuclei of primordial and primary follicles at PND6, as well as in all follicles of PND14 mice treated with GnRH antagonist. By contrast, without CTX exposure, PND14 ovaries undergoing spontaneous HPG stimulation (attested by increased *Cyp19* expression) showed decreased SUMO2/3 in GC of growing medullary follicles. This GC hypo-SUMOylation mirrored a transient TRIM28-S^473P^ immunostaining increase in the same follicles (Supplementary Fig. S3b-g). Altogether, these observations demonstrate that in a physiological context, the HPG axis exerts transient acute negative control on SUMOylation dynamics associated with inactive TRIM28-S^473P^ accumulation.

### Chronic PKA gain-of-function induces hypo-SUMOylation and increased P-TRIM28 from the pre-pubertal stage

Given that TRIM28-SUMO2 conjugates act as gatekeepers of GC identity^5^, we used *Prkar1a* conditional knockout mice (R1α^cKO^: *Prkar1a*^fl/fl^; *Sf1Cre*/+) to explore PKA-TRIM28 interaction consequences on GC identity plasticity. In these mice, developed to study steroidogenic tissue alterations from Carney Complex mutations, conditional *Prkar1a* KO results in PKA-R1α regulatory unit loss and consequent PKA gain-of-function in gonadic somatic cells^58^ (Supplementary Fig. S4a).

R1α^cKO^ female mice were born with normal ovaries (Supplementary Fig. S4b) but were sterile as adults. Histological examination along early postnatal ontogenesis (PND1–5) revealed no striking abnormalities. However, from PND14, atretic secondary follicles increased in the medulla of R1α^cKO^ ovaries compared to R1α^WT^ (*Prkar1a*^fl/fl^), and this was clearly exacerbated in PND30 R1α^cKO^ ovaries (Supplementary Fig. S4b), despite normal FSH and LH plasma concentrations at juvenile (PND21) and pre-pubertal (PND30) stages (Supplementary Fig. S4c).

We then investigated GC plasticity at the key peripubertal PND30 stage. IHC revealed a significant SUMO2/3 immunostaining decrease in GC nuclei across all growing follicles, including peripheral non-atretic follicles, in R1α^cKO^ *versus* R1α^WT^ ovaries (Fig. 4a), confirmed by WB on whole ovary extracts (Supplementary Fig. S4d). This hypo-SUMOylation was associated to increased TRIM28-S^473P^ accumulation (Fig. 4a-b, Supplementary Fig. S4e). TRIM28-SUMO2/3 PLA, analyzing *in situ* SUMO peptide conjugation to TRIM28, revealed significant decreases in both the number of GC nuclei accumulating PLA foci and the number of foci per GC in R1α^cKO^ *versus* R1α^WT^ ovaries (Fig. 4c). Altogether, these data indicate that chronic PKA-GOF drives basal global hypo-SUMOylation, including reduced SUMOylated TRIM28 levels, mirroring the basal increase of inactive TRIM28-S^473P^ accumulation in R1α^cKO^ GC.

**Figure 4:**
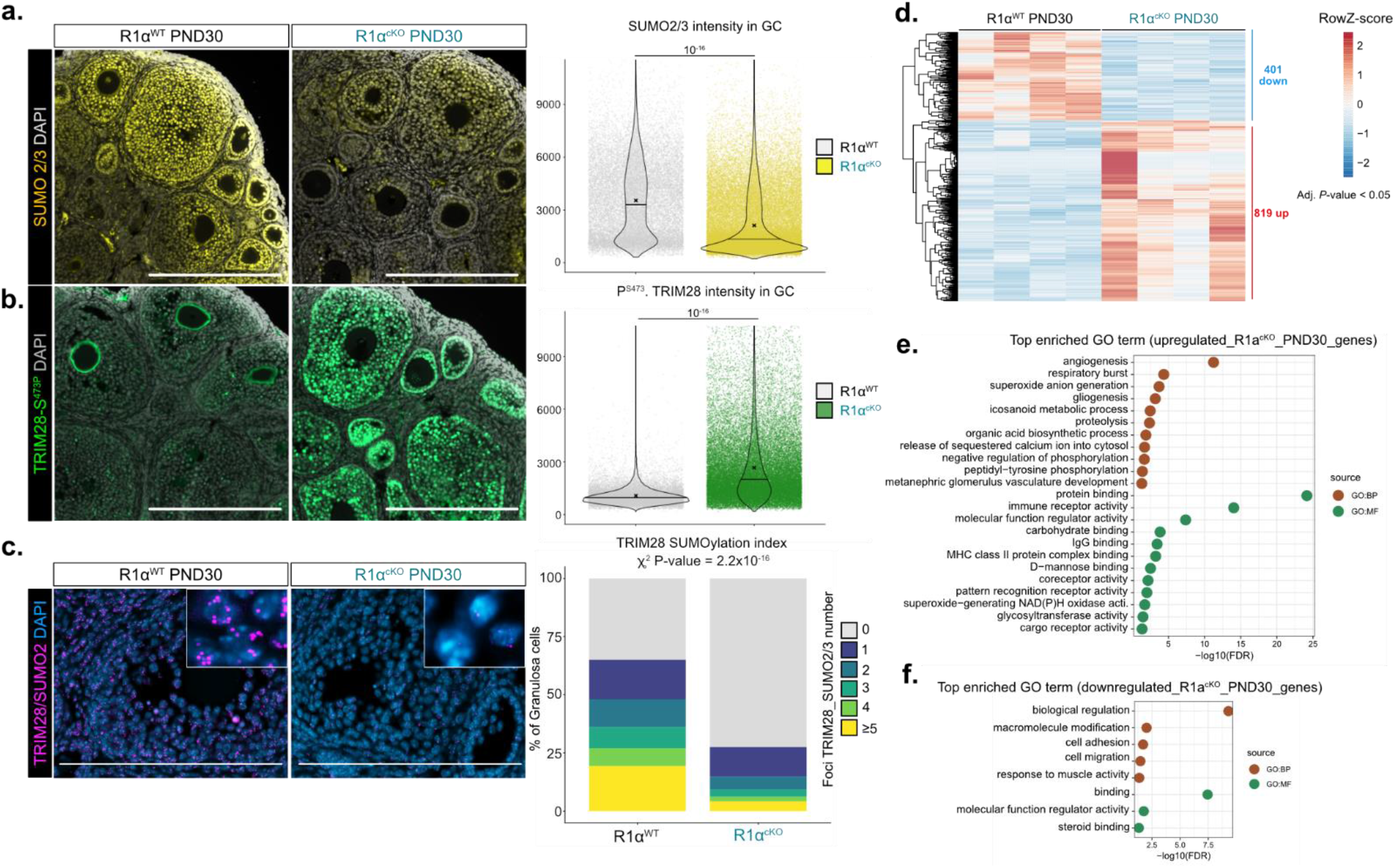
*In vivo* chronic PKA gain of function in ovarian somatic cells induced hypo-SUMOylation associated to the increase of P-TRIM28 levels from the peri pubertal stage. **a.** Immunofluorescence labelling of SUMO2/3 (yellow) in PND30-R1α^WT^ and PND30-R1α^cKO^ ovaries. The graphs quantified SUMO2/3 immunostaining signal intensity in GC nuclei from ovaries of 3/4 mice *per* genotype. *P* value was determined with a Kruskal–Wallis test with multiple comparisons using Dunn’s test. Scale bar: 200µm. **b.** Immunofluorescence labelling of TRIM28-S^473P^ (green) at PND30 R1α^WT^ and R1α^cKO^ ovaries. The graphs quantified TRIM28-S^473P^ immunostaining signal intensity in GC nuclei from ovaries of 3/4 mice *per* genotype. *P* value was determined with a Kruskal–Wallis test with multiple comparisons using Dunn’s test. Scale bar: 200µm. **c.** Proximity ligation assay (PLA) of TRIM28 and SUMO2/3 in GC nuclei of PND30-R1α^WT^ and PND30-R1α^cKO^ ovaries. Histograms represent proportion of GC containing the specified number of dots of interaction per nucleus of R1α^WT^ or R1α^cKO^ ovaries. To assess whether the distribution of cells differs significantly between the groups a χ² test of independence was assessed. n =3 mice *per* genotype. Scale bar: 200µm. **d.** Heatmap representing the median centred expression of dysregulated genes (adj*. P-*value < 0.05) determined with Benjamini and Hochberg method, between R1α^WT^ and R1α^cKO^ 4-week-old ovaries. **e.** Top GO terms from Gene Ontology functional enrichment analysis based on the upregulated genes lists. **f.** Top GO terms from Gene Ontology functional enrichment analysis based on the downregulated genes lists.

To gain insight into gene expression profiles associated with PKA-GOF, we performed comparative bulk RNA-Seq on PND30 R1α^cKO^ (n=5) and R1α^WT^ (n=5) ovaries. Transcriptomic analyses revealed 1220 significantly deregulated genes (819 upregulated, 401 downregulated; *adj.P*<0.05) (Fig. 4d), with most fold changes remaining in a low range (-1<LOG2FC=1 for 613 genes) (Supplementary Fig. S4f, Table Data S1), suggesting limited divergence from R1α^WT^ molecular signatures at this stage. Gene Ontology and GSEA highlighted increased expression in angiogenesis-related genes (*Vcam1*, *Pdgfra*, *Hgs*, *Pecam1*, *Grn*) (Fig. 4e-f) whereas signatures associated with cell homeostasis, such as apoptosis and proliferation, were virtually unaffected (Supplementary Fig. S4g). Altogether, these analyses indicate that chronic ovarian PKA-induced hypo-SUMOylation with increased P-TRIM28 levels has limited consequences on PND30 pre-pubertal somatic cell homeostasis.

### Chronic PKA-GOF redirects adult ovarian somatic cell plasticity toward testicular identity

Given *Trim28*, *Esr1-2*, or *Foxl2* inactivation disrupt ovarian identity from adulthood^3–5,59^, we next explored the 12-week-old R1α^cKO^ ovaries. R1α^cKO^ female mice were sterile from puberty onset. H&E staining at 12 weeks revealed severe gonadal dysplasia: growing follicles at different stages developed into atretic follicles, serous cysts, and pseudo-tubular structures where Sertoli-like cells, characterized by voluminous cytoplasm and large nuclei with three nucleoli, replaced GC (Fig. 5a). IHC confirmed nuclear accumulation of SOX9, DMRT1, and GATA1 in Sertoli-like cells, with concomitant FOXL2 loss. To confirm that ovary failures resulted from PKA overactivation rather than decreased activity, we examined *Sf1Cre*; *Prkar1a*^fl/fl^ *Prkaca*^+/–^ mice with PKA-Cα haploinsufficiency^58^. These mice retained fertility and healthy ovary histology (Supplementary Fig. S5a), establishing that R1α^cKO^ phenotypes stem from PKA overactivation.

**Figure 5:**
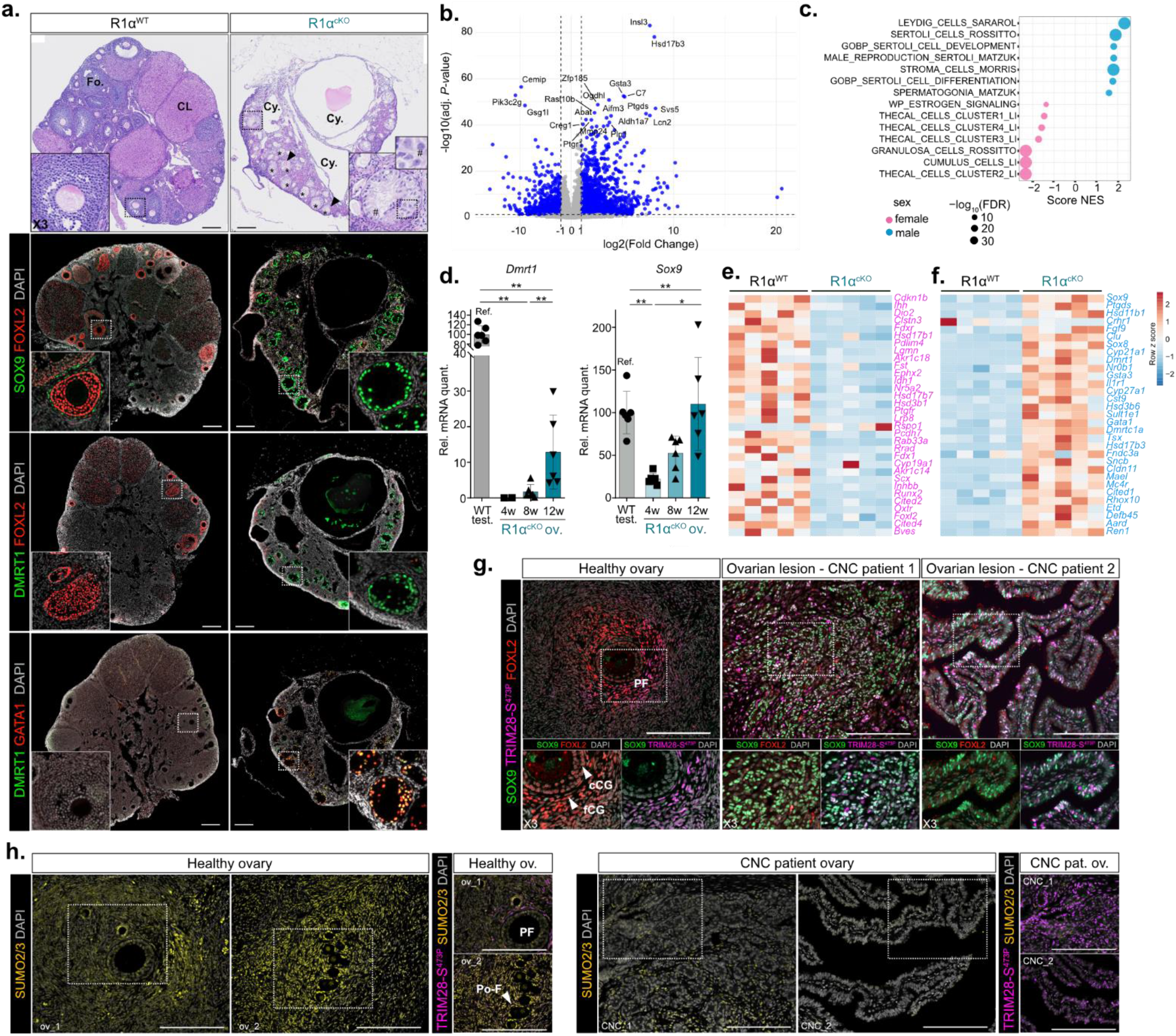
Chronic PKA GOF redirected the adult ovarian somatic cell plasticity. **a.** H&E staining of ovaries from R1α^WT^ and R1α^cKO^ 12-week-old mice. Degenerating oocytes still present in the pseudo tubules are indicated by arrowheads. Inset boxes magnified follicle in R1α^WT^ and pseudo tubular structure containing Sertoli-like cells in R1α^cKO^ ovaries. Immunofluorescence labelling of SOX9 (green), FOXL2 (red), DMRT1 (green) and GATA1 (red) in 12-weeks-old R1α^cKO^ and R1α^WT^ ovaries. *\*: pseudo-tubules, CL: corpus luteum, Fo: follicle, Cy: Cysts, #: Sertoli-like cells.* Scale bars: 200 μm. **b.** Volcano plot showing the distribution of DEG in 12-weeks-old R1α^cKO^ ovaries *vs.* R1α^WT^. **c.** Gene set enrichment analysis (GSEA) of deregulated gene expression data using gene lists associated with granulosa/Sertoli-cell-identity markers and Thecal/Leydig cell markers in 12-weeks-old R1α^cKO^ ovaries compared to R1α^WT^ ovaries^5,105–107^. **d.** RT-qPCR analyses of the *Dmrt1* and *Sox9* transcript levels in 4-, 8- and 12-weeks-old R1α^cKO^ ovaries (*ov*.) *vs.* R1α^WT^ testes (*test*.). Quantification histograms represent mean ± SD. n= 5/6 ovaries *per* genotype. Statistical analysis was performed using two-sited Mann-Whitney U test. \**P*<0.05, \*\**P*<0.01, \*\*\**P* <0.005. **e.** Heatmap representing the median centred expression of significantly downregulation of ovarian expression markers in 12-weeks-old R1α^cKO^ ovaries compared with R1α^WT^. **f.** Heatmap representing the median centred expression of significantly upregulation of testis expression markers in 12-weeks-old R1α^cKO^ ovaries compared to R1α^WT^ ovaries. **g.** Immunofluorescence labelling of FOXL2 (red), TRIM28-S^473P^ (purple) and SOX9 (green) in ovarian lesions of CNC patient (mucinous adenoma-CNC patient 1 and cystic lesion-CNC patient 2) and healthy human ovary sections containing primary follicles (PF) with cubic GC (cGC) and flattened GC (fGC) layers. Scale bars: 200 μm. **h.** Immunofluorescence labelling of SUMO2/3 (yellow) and TRIM28-S^473P^ in ovarian lesions of CNC patient ovaries and healthy human ovaries sections containing primordial follicles (Po-F) and primary follicles (PF). Scale bars: 200 μm.

Unbiased bulk RNA-Seq on 12-week-old R1α^cKO^ (n=5) and R1α^WT^ (n=5) ovaries revealed extensive molecular network deregulation with 7128 DEG (3759 upregulated, 3369 downregulated; *adj.P*<0.05), including 2383 with -1<LOG2FC>1 (Fig. 5b, Supplementary Fig. S5b-e, Table Data S2). GSEA showed robust negative enrichment for granulosa and theca cell signatures and estrogen signaling, opposed by positive enrichment for Sertoli and Leydig cell gene sets (Fig. 5c). RT-qPCR confirmed significant *Sox9* mRNA increase between 4 and 12 weeks, with delayed *Dmrt1* upregulation (Fig. 5d). Heatmaps of 64 gonadic cell identity markers and GO analyses demonstrated repression of follicle plasticity and GC identity modulators (*Foxl2, Lrh1/Nr5a2*, *Runx2*, *Ihh*, *Cited4*, *Hsd17b7*, *Fst*, *Hsd3b1*, *Hsd17b7*) and gain of testicular fate markers (*Dmrt1*, *Sox9*, *Ptgds*, *Fgf9*, *Gata1*, *Cldn11*, *Insl3*, *Hsd17b3*, *Hsd3b6*) in adult R1α^cKO^ ovaries (Fig. 5e-f, Supplementary Fig. S5d-h).

Hormonal assays revealed a 41.7-fold testosterone increase in 12-week-old R1α^cKO^ ovaries (3.34 ng/organ *vs*. 0.08 ng in R1α^WT^) (Supplementary Fig. S6a). Plasma testosterone also increased significantly (0.45 ng/mL *vs*. 0.14 ng/mL in R1α^W^), while levels remained unchanged until PND17 (Supplementary Fig. S6b). This androgen excess associated with chronic upregulation of *Cyp17*, *Hsd17b3* and *Hsd3b6* mRNA (Supplementary Fig. S6c-e), despite low LH plasma levels (Supplementary Fig. S6f) and unaffected cholesterol metabolism (Supplementary Fig. S6g). IHC revealed an increase in stromal cells expressing CYP17, LHCGR associated to chronic PKA GOF, while VCAM-1 (Supplementary Fig. S6h), demonstrated the impact of androgen overproduction on the stroma^60^.

To dissociate PKA cell-autonomous effects from androgen signaling, we analyzed *Prkar1a*^fl/fl^; *Ar*^fl/fl^; *Sf1Cre*/+ (Ar-R1α^dKO^) mice. Comparative HE and IHC indicated pseudo-tubular structures with nuclear SOX9 and DMRT1 accumulation in 12-week-old Ar-R1α^dKO^ ovaries, similar to R1α^cKO^ (Supplementary Fig. S7a). RT-qPCR and hormonal assays confirmed similar *Sox9*, *Dmrt1*, and *Hsd17b3* upregulation with increased testosterone in both Ar-R1α^dKO^ and R1α^cKO^ ovaries (Supplementary Fig. S7b-e), establishing that chronic PKA-GOF disrupts ovarian somatic cell plasticity independently of androgen signaling.

### PRKAR1A inactivating mutation promotes hypo-SUMOylation in CNC ovarian lesions with P-TRIM28 and SOX9 accumulation

CNC is a rare autosomal dominant multiple neoplasia syndrome caused by inactivating mutations in *PRKAR1A*^61,62^, resulting in chronic PKA activation — the precise molecular context modeled by our R1α^cKO^ mice. CNC women frequently develop serous cystic lesions and occasionally ovarian mucinous adenocarcinoma^63^. IHC on biopsies from two CNC patients revealed that in cystic lesions (patient 2) and mucinous adenoma (patient 1), cells were devoid of nuclear SUMO2-conjugated immunostaining but strongly accumulated TRIM28-S^473P^ (Fig. 5g-h, Supplementary Fig. S8a), unlike healthy stromal cells and flattened GC surrounding oocytes in control follicles. Co-immunostaining indicated that hypo-SUMOylated CNC lesion cells accumulated both TRIM28-S^473P^ and SOX9, and to a lesser extent DMRT1, in their nuclei. RT-qPCR on a third independent CNC ovary biopsy revealed decreased *Trim28*, *Pias3* and *Sumo2* transcripts with a dramatic *Sox9* increase (Supplementary Fig. S8b). Thus, *PRKAR1A* inactivating mutation-driven chronic PKA activity in CNC downregulates SUMOylation processes and TRIM28 activity, a major component of ovarian somatic cell identity, and promotes emergence of testicular signatures, mirroring the R1α^cKO^ mouse model.

### PKA-induced TRIM28 phosphorylation inhibits chromatin binding

Adult R1α^cKO^ and Trim28^cKO^ ovaries share similar histological features with pseudo-tubular structures hosting Sertoli-like cells from puberty^5^. Comparing DEG (*adj.P*<0.05, - 1<LOG2FC>1) revealed 1242 genes (52%) commonly deregulated in both models in the same direction (Fig. 6a, Supplementary Fig. S9a, Table Data S3), including networks involved in development, tissue differentiation, and homeostasis (Supplementary Fig. S9b). These analyses confirmed that converging histologies relied on common drastic expression changes in gonadal somatic cell plasticity genes (*Dmrt1*, *Fgf9*, *Ptgds*, *Gata1*, *Sox8*, *Lrh1/Nr5a2*, *Fst*, *Ihh*, *Cited4*, *Hsd17b7*, *Hsd3b1*, *Prdm6*) (Fig. 6a).

**Figure 6:**
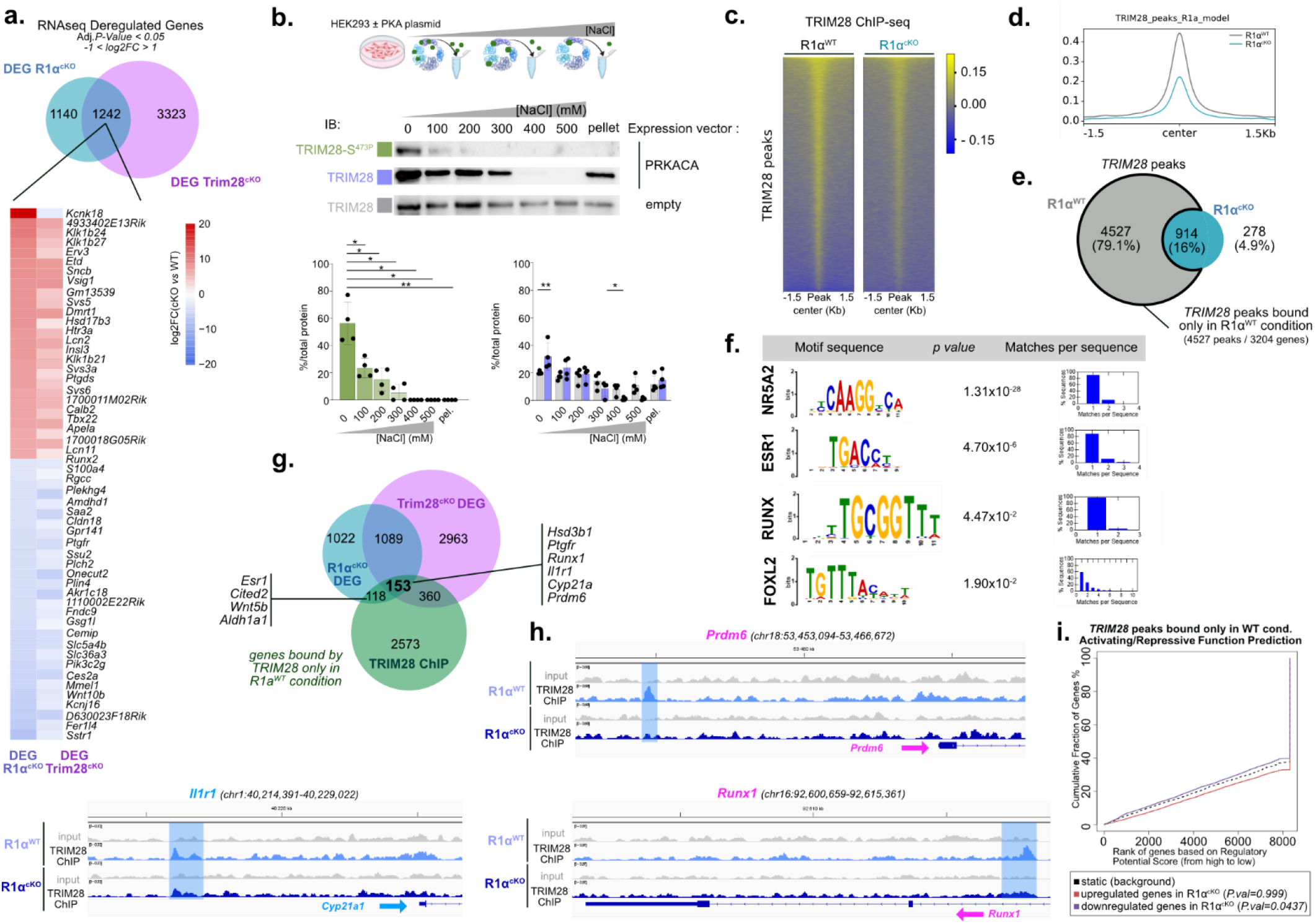
PKA-induced TRIM28 chromatin binding inhibition alters expression of TRIM28 target genes involved in gonadic somatic cell identity. **a.** Venn diagram of commonly deregulated genes in adult R1α^cKO^ and Trim28^cKO^ ovaries. Bottom panel: heatmap of the top 50 most commonly DEG in both models. **b.** TRIM28-S^473P^ chromatin binding affinity measurement. Upper panel: Schematic design of chromatin preparation and sequential salt extraction. Chromatin isolated from HEK293T cells transfected with PKA-Cα or empty vector was extracted with increasing salt concentrations. Bottom panel: WB using anti-TRIM28-S^473P^ and anti-TRIM28 antibodies. PCNA immunostaining used as loading/extraction control in Supplementary Fig. S9. Quantification histograms represent mean ± SEM. n=4 independent experiments. Statistical analysis was performed using two-way ANOVA followed by Holm-Šídák’s multiple comparisons test. *P<0.05, **P<0.01, ***P<0.005, ****P<0.0001. **c.** Heatmap of normalized TRIM28 ChIP-seq reads around TRIM28 peak center in 12-week-old R1α^WT^ *vs* R1α^cKO^ ovaries. Yellow traces represent TRIM28 ChIP-seq reads (± 1.5kb from the peak center). **d.** Metaprofile of average TRIM28 signal around the TRIM28 peak centers (±1.5 kb) in adult R1α^WT^ and R1α^cKO^ ovaries. **e.** Venn diagram showing that 4,534 of 5,448 TRIM28 peaks (79.1%) are R1α^WT^ specific (*ie*. lost in R1α^cKO^). **f.** Binding motifs enrichment for TFs involved in GC identity based on TRIM28 ChIP-seq sequences that were specific to R1α^WT^ ovaries. **g.** Venn diagram comparing genes for which TRIM28 peaks were specific to R1α^WT^ ovaries with the DEG in R1α^cKO^ and Trim28^cKO^ ovaries. **h.** Example of signal intensity tracks showing TRIM28 peaks lost in/around genes in R1α^cKO^, with upregulated (blue) or downregulated (pink) expression genes in R1α^cKO^. **i.** Prediction of active/repressive function associated to TRIM28 peaks specific to R1α^WT^ chromatin. Distribution of genes based on their deregulation level (upregulated in R1α^cKO^ ovaries in blue and downregulated in red) and binding by TRIM28. Genes are classified as potential regulatory targets with a *regulatory potential score* based on the presence of a TRIM28 peak and the degree of their deregulation. The *P* values indicate the statistical significance of the distributions of the groups of upregulated or downregulated genes compared to the group of genes that are not differentially expressed.

To test whether PKA-induced phosphorylation alters TRIM28 chromatin engagement, we performed sequential salt extraction of chromatin-bound proteins from H293 cells after PKA-Cα ectopic expression. Under basal conditions, endogenous TRIM28 was recovered across all fractions, including the final pellet (Fig. 6b, Supplementary Fig. S9c). Upon PKA activation, TRIM28 shifted toward less tightly bound fractions with marked reduction in the 400 and 500 mM NaCl eluates, remaining only in the pellet fraction. This suggests a small TRIM28 fraction remained either inaccessible to PKA-mediated phosphorylation or not displaced by phosphorylation alone. Importantly, TRIM28-S^473P^ was specifically excluded from high-affinity fractions and recovered in lower-salt fractions, indicating PKA-dependent phosphorylation is incompatible with strong chromatin retention (Fig. 6b). Moreover, given TRIM28-S^473P^ is detected only in the 0 mM NaCl fraction, and not in the other low-affinity fractions in which TRIM28 is present, could reflect phosphorylation at Ser489 present in those fractions, which would be consistent with the sequential phosphorylation model observed *in vitro*. To directly address Ser473 and Ser489 contributions, we performed sequential salt extraction after transient transfection of WT-TRIM28 or phosphomimetic mutants (S473E, S489D, S473E/S489D). While WT-TRIM28 retained robust chromatin association, both single mutants showed markedly reduced ability to remain in deeply bound fractions, including the insoluble pellet, with no additive effect in the double mutant (Supplementary Fig. S9d). Thus, phosphorylation at either residue suffices to destabilize chromatin anchoring.

Next, we explored PKA-GOF impact on endogenous TRIM28 genome-wide chromatin binding by ChIP-Seq in R1α^WT^ and R1α^cKO^ ovaries (biological duplicates). We identified 5441 TRIM28 binding sites in R1α^WT^ chromatin *versus* 1192 in R1α^cKO^, with 914 common, indicating 4527 sites (79%) were lost in R1α^cKO^ (Fig. 6c-d, Table Data S4-5). These 4527 sites mapped to 3204 closest genes (Fig. 6e, Table Data S6). Genomic mapping revealed distribution into intergenic (45.72%), intronic (39.74%), and promoter regions (10.15%) (Supplementary Fig. S9e). Motif enrichment analysis showed that TRIM28-bound sites in R1α^WT^ (lost in R1α^cKO^) were co-enriched with LRH1/NR5A2, ESR1/2, RUNX, and FOXL2 motifs (Fig. 6f), suggesting functional interaction with these key ovarian factors. To identify TRIM28 transcriptional targets regulated by PKA-GOF, we integrated RNA-seq and ChIP-seq data, finding 271 deregulated genes in R1α^cKO^ ovaries containing at least one lost TRIM28 binding site. These included ovarian signature genes (*Edn2*, *Prdm6*, *Ephx2*, *Runx1*, *Hsd3b1*) and testicular signature genes (*Hsd17b3*, *Cldn11*, *Il1r1*) (Fig. 6g, Table Data S7). Examples included *Runx1* and *Prdm6* (Fig. 6h, Supplementary Fig. S9f), which are expressed in GC^64,65^, and *Il1r1* related to testicular signature^5^. BETA analysis (peaks within ±100 kb of TSS)^50^ showed that TRIM28 binding sites lost in R1α^cKO^ were associated with downregulated genes, supporting TRIM28 as a transcriptional co-activator in R1α^WT^ ovaries (Fig. 6i). Together, these results reveal that chronic PKA activity inhibits TRIM28 chromatin binding, impacting expression of target genes involved in somatic cell identity.

### Chronic PKA-GOF destabilizes FOXL2-TRIM28 complexes and their chromatin binding

TRIM28 is an integrative chromatin protein that does not bind DNA directly but is recruited to chromatin close to FOXL2 to maintain ovarian pathways and repress testicular genes^5^. We assessed TRIM28-FOXL2 interactions in FOXL2-positive GCs of remaining follicles in R1α^cKO^ ovaries at PND30 and 12 weeks using PLA. Analyses revealed significant decreases in TRIM28-FOXL2 interactions in GC nuclei from PND30, worsening by 12 weeks in R1α^cKO^ *vs* R1α^WT^ (Fig. 7a-b, Supplementary Fig. S10a), highlighting early prepubertal complex destabilization by chronic PKA-GOF. Consistent with these observations, PLA in WT ovaries during gonadotropic kinetics showed significant transient decreases in GC nuclei accumulating TRIM28-FOXL2 foci and in foci number *per* GC after 2 hours of eCG and 2–4 hours of hCG (Supplementary Fig. S10b), indicating gonadotropic hormone cyclicity exerts transient negative control on complex stability. Next, we performed FOXL2 ChIP-Seq in 12-week-old R1α^cKO^ and R1α^WT^ ovaries. Comparison of heatmaps identified 50864 FOXL2 binding sites in R1α^WT^ *versus* 44114 in R1α^cKO^, with 27852 common, indicating 23012 sites (34.3%) were no longer occupied in R1α^cKO^ (Fig. 7c-d, Supplementary Fig. S10c, Table Data S8-9). These mapped to 9176 closest genes (Fig. 7d-e, Table Data S10). Genomic mapping revealed distribution into intergenic (45.4%), intronic (48.73%), and promoter regions (2.49%) (Supplementary Fig. S10d). Motif enrichment showed FOXL2-bound sites in R1α^WT^ (unoccupied in R1α^cKO^) were co-enriched with LRH1/NR5A2, ESR1, ESR2, and FOXL2 motifs (Fig. 7f).

**Figure 7:**
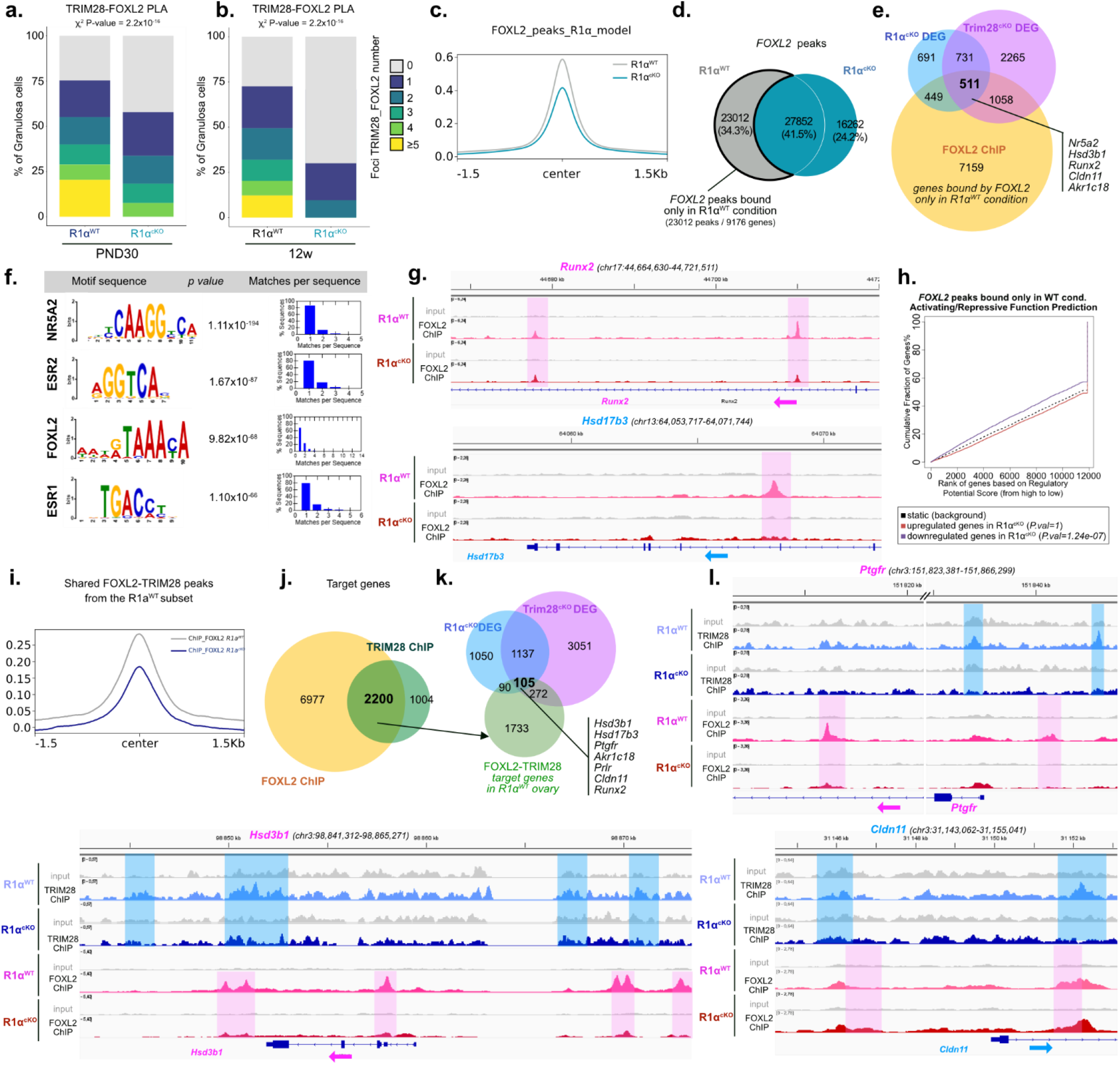
*In* vivo chronic PKA GOF destabilizes FOXL2-TRIM28 complexes in GC. **a. b.** PLA analyses of TRIM28-FOXL2 interactions in GC nuclei of R1α^cKO^ and R1α^WT^ ovaries (PND30 and 12-week-old). Proportion of GC nuclei containing the number of TRIM28– FOXL2 PLA dots *per* nucleus. n=3 ovaries *per* genotype/age. Corresponding images in Supplementary Fig. S10a. **c.** Metaprofile of average FOXL2 signal around the FOXL2 peak center in adult R1α^WT^ and R1α^cKO^ ovaries. **d.** Venn diagram showing that 34.5% of FOXL2 peaks are R1α^WT^ specific. **e.** Venn diagram comparing genes for which FOXL2 peaks were specific to the 12-week-old R1α^WT^ ovaries with the DEG in R1α^cKO^ and Trim28^cKO^ ovaries. **f.** Enrichment of binding motifs for TFs involved in GC identity based on FOXL2 ChIP-seq sequences that were specific to 12-week-old R1α^WT^ ovaries. **g.** Example of FOXL2 peaks that were lost in/around genes in the R1α^cKO^ ovaries and expression of which was altered in R1α^cKO^ models. **h.** Prediction of active/repressive function associated to FOXL2 peaks specifically present in R1α^WT^ chromatin. Distribution of genes based on their deregulation level (upregulated in R1α^cKO^ ovaries in blue and downregulated in red) and their binding by FOXL2. Genes are classified as potential regulatory targets with a *regulatory potential score* based on the presence of a FOXL2 peak and the degree of their deregulation. *P* values validate distribution significance compared DEG groups to non-DEGs. **i.** Metaprofile of FOXL2 signal average around the FOXL2 peak centers from the R1α^WT^ specific TRIM28 peaks (±1.5kb) in R1α^WT^ and R1α^cKO^ adult ovaries. **j.** Venn diagram comparing genes for which both FOXL2 and TRIM28 peaks were lost in R1α^cKO^ ovaries. **k.** Venn diagram comparing 2200 genes for which both FOXL2 and TRIM28 peaks were lost in R1α^cKO^ ovaries with DEG in R1α^cKO^ and Trim28^cKO^ ovaries. **l.** Example of FOXL2 and TRIM28 peaks lost in/around genes in R1α^cKO^, with upregulated (blue) or downregulated (pink) expression genes in R1α^cKO^.

In R1α^cKO^ ovaries, 960 deregulated genes contained at least one *cis*-binding site no longer occupied by FOXL2, including downregulated ovarian genes (*Lrh1/Nr5a2*, *Prdm6*, *Ephx2*, *Runx2*, *Hsd3b1*, *Edn2*, *Fst*) and upregulated testicular genes (*Hsd17b3*, *Fgf9*, *Dmrt1*, *Cldn11*, *Il1r1*), plus *Wt1* and *Esr1* (Fig. 7g, Supplementary Fig. S10e, Table Data S11). BETA analysis showed FOXL2 binding sites lost in R1α^cKO^ strongly associated with downregulated genes, supporting FOXL2 acted primarily as transcriptional activator rather than repressor in R1α^WT^ ovaries (Fig. 7h). This suggested direct FOXL2 involvement in PKA target gene regulation, potentially with TRIM28. Heatmap analyses identified a first group of 1004 genes with TRIM28 but not FOXL2 motifs, and a second group of 2200 genes with both FOXL2 and TRIM28 motifs were no longer occupied in R1α^cKO^ (Fig. 7i-j, Supplementary Fig. S11a). Among genes displaying both loss of TRIM28-FOXL2 co-bindings and strong deregulated expression in R1α^cKO^ (-1<LOG2FC>1, *adj.P*<0.05), we identified 195 genes (Fig. 7j-k, Table Data S12), exemplified in Figure 7l and Supplementary Fig. S11eb. Notably, 105 TRIM28-FOXL2 co-bound genes were deregulated in both R1α^cKO^ and Trim28^cKO^ (57% upregulated, 33% downregulated) (Supplementary Fig. S11c-d, Table Data S12). Again, this list included genes controlling follicle plasticity, GC differentiation, and steroidogenesis (*Kitl*, *Aldh1a3*, *Lifr*, *Prlr*, *Edn2*, *Runx2*, *Pparg*, *Prdm6*, *Akr1c18*, *Ptgfr*)^65–74^, plus Sertoli cell signature genes (*Cldn11, Gja6, Wipf3, Kcng4* )^58^ and testicular homeostasis regulators (*Il1r1*, *Hsd17b3* )^75–77^ (Fig. 7l, Supplementary Fig. S11b). Thus, chronic PKA activity impairs ovarian somatic cell plasticity by destabilizing FOXL2-TRIM28 chromatin interactions that drive identity-maintaining genetic programs.

### Chronic PKA activity exerts TRIM28-independent controls disrupting ovarian identity maintenance

The incomplete DEG overlap between adult R1α^cKO^ and Trim28^cKO^ transcriptomes suggested chronic PKA activity exerts additional control over ovarian identity beyond direct TRIM28 inhibition. We therefore monitored trans-differentiation in *Trim28*^fl/fl^; *Prkar1a*^fl/fl^; *Sf1Cre*/+ (R1α-T28^dKO^) mice. As with R1α^cKO^ females, R1α-T28^dKO^ mice showed no embryonic gonad abnormalities or ovarian weight differences up to PND14. However, from PND27, cysts and pseudo-tube structures began to develop, and by 5 weeks, a large number had invaded the entire ovary, indicating extensive earlier cortical and medullary follicle trans-differentiation at puberty onset (Fig. 8a, Supplementary Fig. S12a-b). IHC confirmed nuclear SOX9 and DMRT1 accumulation with concurrent FOXL2 and SUMO2/3 loss in pseudo-tube cells (Supplementary Fig. S12c), suggesting synergistic effects of TRIM28 loss and PKA-GOF on ovarian identity. To understand the molecular basis, we compared 5-week-old R1α-T28^dKO^, R1α^cKO^, Trim28^cKO^, and R1α^WT^ ovarian transcriptomes (n=4 each) by bulk RNA-Seq. PCA revealed clear separation of R1α-T28^dKO^ transcriptomes from Trim28^cKO^ and R1α^cKO^, the latter one remained close to R1α^WT^ (Supplementary Fig. S13a). DEG numbers increased from 605 in PND30 R1α^cKO^ to 2583 in 5-week-old R1α^cKO^, confirming the peripubertal period importance for establishing adult ovarian identity. R1α-T28^dKO^ showed amplified deregulation with 4117 DEG (3124 upregulated, 994 downregulated), including 1671 (40%) specific to R1α-T28^dKO^ and 717 (27.7%) common to all three mutants (Fig. 8b-c, Table Data S13). GO and GSEA of R1α-T28^dKO^ transcriptomes revealed strong positive enrichment for Sertoli cell differentiation and, to a lesser extent, Leydig cell differentiation and androgen signaling, mirrored by negative enrichment for female sexual characteristic development, follicle development, granulosa/theca cells, and estrogen signaling (Fig. 8d, Supplementary Fig. S13b-c). The specific R1α-T28^dKO^ DEG cluster showed strong downregulation of key ovarian identity genes (*Foxl2*, *Esr2*) and theca cell differentiation/follicle maturation genes (*Ihh*, *Fshr*, *Fst*, *Inha*, *Cyp19*), attesting to more pronounced, accelerated identity loss (Fig. 8c, Table Data S14). Surprisingly in this cluster, we also identified marked *Amh* downregulation (Table Data S14). IHC confirmed AMH loss in R1α-T28^dKO^ ovaries (Supplementary Fig. S13f). Given previous studies demonstrated AMH protective role for cortical follicles^78^, the R1α-T28^dKO^genetic context inducing pronounced AMH loss could mobilize this reserve follicle activation toward earlier trans-differentiation in R1α-T28^dKO^, and would explain the explosive pseudo-tube increase at 5 weeks. Synergy between PKA-GOF and TRIM28 loss also impacted shared DEG clusters, with downregulation of ovarian genes such as *Nr5a2*, *Inhba*, *Lhcgr*, *Prlr*, and *Bhmt* (Fig. 8c, Supplementary Fig. S13d-e).

**Figure 8:**
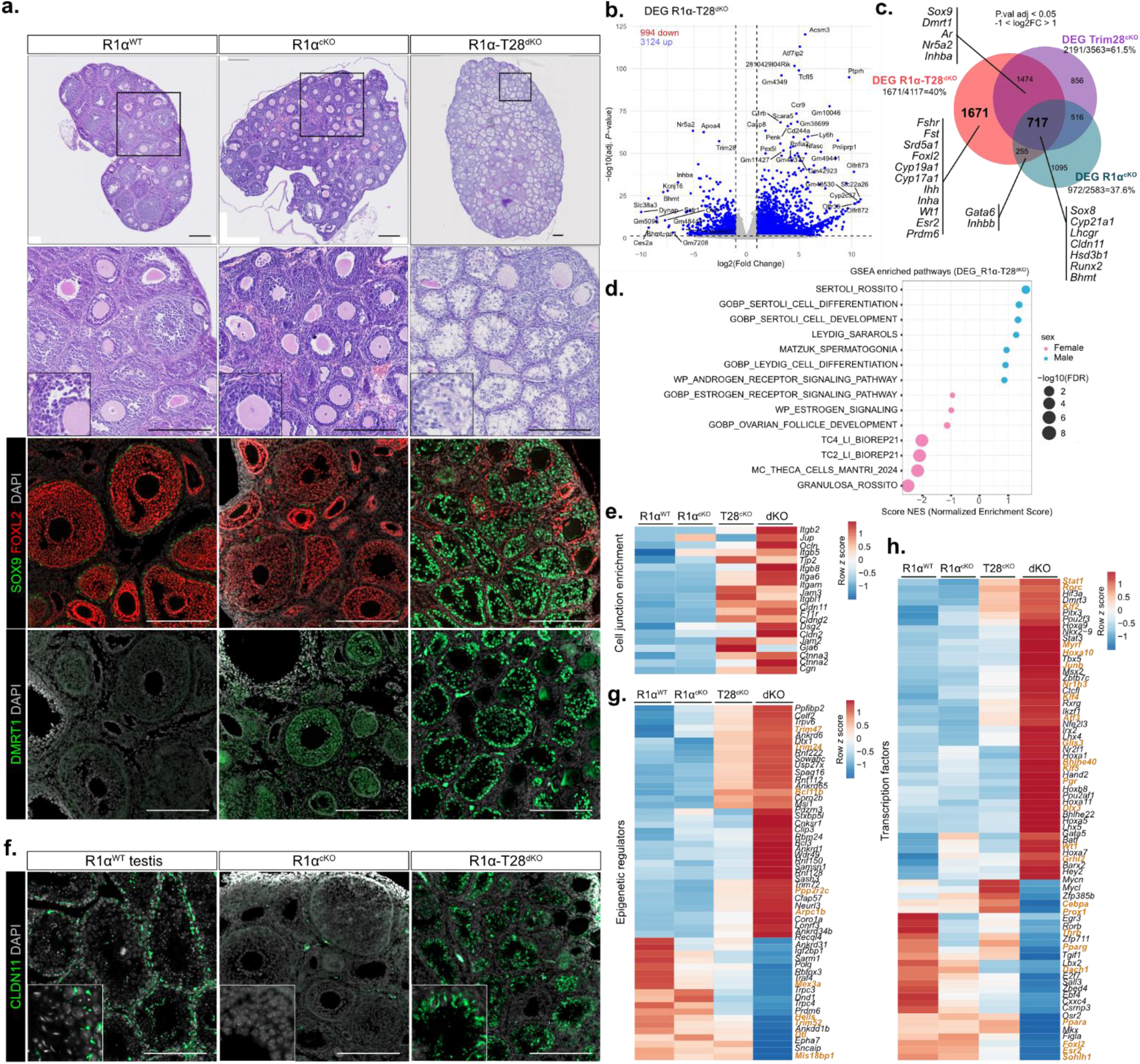
Chronic PKA activity exerts TRIM28-independent controls disrupting ovarian identity maintenance. **a.** H&E staining of R1α^WT^, R1α^cKO^ and R1α-T28^dKO^ 5-week-old ovaries. Immunofluorescence labelling of SOX9 (green), FOXL2 (red) and DMRT1 (green) in R1α^WT^, R1α^cKO^ and R1α-T28^dKO^ 5-week-old ovaries. Scale bars: 200 μm. **b.** Volcano plot showing the distribution of deregulated genes in 5-weeks-old R1α-T28^dKO^ ovaries *vs* R1α^WT^ ovaries. **c.** Venn diagram illustrating the commonly deregulated expressed genes (-1<LOG2FC>1, adj*. P-value* <0.05) in both R1α^cKO^, Trim28^cKO^ and R1α-T28^dKO^ 5-week-old ovaries compared to R1α^WT^ ovaries. **d.** Gene set enrichment analysis (GSEA) of deregulated gene expression data using gene lists associated with granulosa/Sertoli-cell-identity markers in 5-week-old R1α-T28^dKO^ ovaries compared to the R1α^WT^ ovaries. **e.** Heatmap representing the median centred expression of significantly upregulated genes encoding Sertoli cell junction markers in 5-week-old R1α-T28^dKO^ ovaries compared to R1α^cKO^, Trim28^cKO^ and R1α^WT^ ovaries. **f.** Immunofluorescence labelling of CLDN11 (green) in 5-week-old, R1α^cKO^ - and R1α-T28^dKO^ ovaries and 8-week-old R1α^cKO^ testis. Scale bars: 200 μm. **g.** Heatmap representing the median centred expression of significantly deregulated genes encoding epigenetic regulators in 5-week-old R1α-T28^dKO^ ovaries compared to R1α^cKO^, Trim28^cKO^ and R1α^WT^ ovaries. Epigenetic regulators for which related to SUMOylation processes has been reported in the literature are shown in orange bold (related references are listed in Table S6). **h.** Heatmap representing the median centred expression of significantly deregulated genes encoding TFs in 5-week-old R1α-T28^dKO^ ovaries compared to R1α^cKO^, Trim28^cKO^ and R1α^WT^ ovaries. Factors for which SUMOylation has been reported in the literature are shown in orange bold (related references are listed in Table S6).

Testicular somatic cell identity signatures in shared DEG clusters showed increased upregulation in R1α-T28^dKO^, including Sertoli cell genes (*Dmrt1*, *Sox9*, *Sox8*, *Wt1*, *Rarres1*, *Etd*, *Cldn11*, *Clu*) while key steroidogenic genes remained downregulated (*Cyp11a1*, *Hsd3b1*, *Cyp17a1*) (Fig. 8c, Supplementary Fig. S13d-e). Furthermore Increased Sertoli cell junction gene expression attested to more pronounced GC-to-pseudo-Sertoli trans-differentiation in R1α-T28^dKO^, confirmed by IHC showing Claudin-11 membrane localization (Fig. 8e-f).

Finally, we extracted expression levels of 690 epigenetic regulation genes^79^ and 1374 TF genes^80,81^, finding strong deregulation specific to R1α-T28^dKO^ for 55 epigenetic regulator and 72 TF genes, including SUMOylation targets, and SUMOylation-related regulators (Fig. 8g-h, Supplementary Fig. S13g, Table S6). Thus, beyond direct TRIM28 inhibition, PKA-GOF targets additional SUMOylation-connected pathways associated with accelerated ovarian identity disruption.

## DISCUSSION

Whereas PKA mediates gonadotrophin effects, its role in maintaining postnatal granulosa cell identity had been unexplored. Here, we show the PGH-cAMP/PKA axis exerts rapid, transient, coordinated negative control over SUMO2/3-conjugation in mouse ovary *in vivo*. Applying HPG axis synchronization followed by gonadotropic stimulations^45^, we highlighted follicular SUMO2/3 hypo-SUMOylation waves recurring transiently in response to consecutive hormonal signals. This transience is consistent with studies showing that SUMO2/3-driven global SUMOylation preserves oocyte development at folliculogenesis end^82^, and that its disruption affects folliculogenesis regulatory processes, including oocyte-to-granulosa communication^83^. Post-translational modifications as core follicular development regulators^84^ further underscores our SUMOylation dynamic relevance. These observations extend to the ovary, our previous adrenal cortex findings that ACTH/PKA signaling acutely downregulates nuclear SUMOylation *via* coordinated SUMO machinery transcriptional reprogramming^16,33^. The restricted transcriptional modulation of SUMO players in primary GC *vs* whole-ovary suggests stromal/theca cells broaden gonadotrophin responses *in vivo*, or that forskolin/IBMX exposures do not fully recapitulate FSH/LH receptor signaling complexity. Notably, hypo-SUMOylation wave rapidity implies post-transcriptional and post-translational mechanisms, including direct PKA phosphorylation of SUMO players, contribute to acute SUMO conjugation dynamics. This dual model echoes multi-layered SUMOylation regulation in adipocyte differentiation^17^.

A central finding is TRIM28 as a direct PKA substrate with functional consequences on its E3-SUMO ligase activity. We identified Ser473 and Ser489 as direct PKA targets and as the two sites with most robust PKA-dependent phosphorylation. Ser473 was previously established as an ATM-dependent site critical for DNA damage responses, where its phosphorylation disrupts HP1 binding and relieves TRIM28-mediated chromatin compaction^52,53^. Our study shows this site also responds to PKA signaling, suggesting Ser473 functions as a convergent phosphorylation hub integrating distinct kinase inputs to modulate TRIM28 chromatin functions. More remarkably, we identified Ser489 as a novel PKA-responsive phospho-site within the HP1-binding domain, overlapping the HP1 box motif. Ser489 position within this interface, provides another structural feature for its phosphorylation functional consequences by altering HP1 recognition and the TRIM28–HP1 heterochromatin complex stability.

*In vitro* kinase assays specified that Ser473 phosphorylation precedes and facilitates Ser489 phosphorylation, implying a conformational relay where Ser473 modification may relieve intramolecular constraints or expose Ser489 to PKA. Sequential phosphorylation cascades are documented for other chromatin proteins, including progesterone receptor where serine-phosphorylation blocks lysine-SUMOylation to derepress transcription^32^, illustrating phosphorylation–SUMOylation crosstalk as a hierarchical TF regulator^31^.

The *in vivo* relevance of the PKA–TRIM28 axis was established through complementary approaches. First, transient TRIM28-S⁴⁷³^P^ accumulation increase in GC nuclei, mirroring SUMO2/3 loss kinetics after PGH exposures confirmed the *in vitro* mechanism operates in the ovarian folliculogenesis. Furthermore, PLA revealed transient interactions between PKA-Cα and TRIM28 in GC nuclei, rapidly reaching a short-lived peak following the administration of each of the gonadotropic exposures thereby confirming the existence of cyclic and reversible interactions. Exploitation of the postnatal mini-puberty window^55^ provided additional physiological relevance: at PND14, spontaneous gonadotrophin surges drive transient ovarian response^56,57^, and we observed spontaneous GC SUMO2/3 staining decrease with increased TRIM28-S⁴⁷³^P^ accumulation in growing medullary follicles, both abolished by GnRH antagonist pretreatment. This demonstrates that PKA–SUMOylation coupling is not a pharmacological artifact but operates during endogenous hormonal pulses, and it reveals the immature ovary is already subjected to HPG axis-driven SUMOylation dynamics at follicular reserve establishment.

The R1α^cKO^ mouse model, where PKA regulatory subunit R1α loss creates constitutive PKA– GOF in gonadic somatic cells^58^, proved instrumental for dissecting chronic PKA–TRIM28 interaction consequences. The R1α^cKO^-ovary progression—from normal ovaries at birth and juvenile stages, with limited transcriptomic perturbation, to pronounced trans-differentiation at adulthood—reveals the ovarian identity-maintenance network is resilient to chronic PKA activation during prepuberty but becomes vulnerable once full cyclical HPG axis output is running after puberty. The transcriptomic trajectory of R1α^cKO^ ovaries closely parallels histopathological progression: peripubertal transcriptome indicates minimal cell identity program disruption, while adult transcriptome showed sex-specific somatic cell signature disruption, with robust granulosa/theca cell and estrogen signaling gene set loss and Sertoli/Leydig cell program gain. SOX9, DMRT1, and GATA1 accumulation in Sertoli-like cells with concomitant FOXL2 loss phenocopies ovarian-to-testicular trans-differentiation described upon *Foxl2* ablation in adult mice^4^ and *Trim28* deletion in GC^5^, firmly placing PKA– TRIM28–SUMOylation axis within the established adult ovarian identity maintenance regulatory network. The significance of shared DEG between R1α^cKO^ and Trim28^cKO^ adult ovaries further supports that substantial PKA-GOF transcriptomic effects are TRIM28-mediated.

Using Ar-R1α^dKO^ mice, lacking PKA-R1α and androgen receptor, we observed ovarian trans-differentiation in R1α^cKO^ mice occurs independently of androgen signaling. This dissociates cell-autonomous PKA-GOF consequences on somatic cell fate from androgen excess effects, previously established as ovarian stromal remodeling contributor^60^. Although R1α^cKO^ ovaries produced excessive testosterone, likely through chronic *Cyp17*, *Hsd17b3*, and *Hsd3b6* upregulation, the trans-differentiation process itself is driven by PKA–TRIM28 mechanism rather than androgen-mediated positive feedback. This distinction has implications for androgen excess disorders like PCOS, where elevated androgens contribute to upset the pituitary and ovarian endocrine activities, but may not initiate the GC identity interruption.

The extension to human pathology through Carney Complex patient ovarian biopsy analysis provides compelling translational validation. CNC ovarian lesions displayed hypo-SUMOylated cells accumulating TRIM28-S⁴⁷³^P^, SOX9, and DMRT1, recapitulating R1α^cKO^ ovary molecular characteristics. RT-qPCR showing decreased TRIM28, PIAS3, and SUMO2 transcripts alongside dramatic SOX9 increase further supports PKA–TRIM28–SUMOylation axis disruption in CNC lesions. Although clinical observations rely on few samples due to CNC ovarian tissue biopsy scarcity, these findings remain significant because CNC ovarian lesions are clinically classified as predominantly serous cystic lesions with mucinous adenocarcinoma predisposition^85^. Our data suggest at least a subset of CNC ovarian lesions may arise from somatic cells that have lost critical ovarian identity genes through chronic PKA-driven TRIM28 inactivation and hypo-SUMOylation, potentially explaining histological diversity.

Chromatin fractionation experiments provided a mechanistic bridge between biochemical PKA–TRIM28 interaction and *in vivo* genomic consequences. Salt extraction assays demonstrated PKA activation shifts TRIM28 toward less tightly chromatin-bound fractions, with TRIM28-S⁴⁷³^P^ specifically excluded from high-affinity chromatin fractions. Phosphomimetic mutagenesis confirmed either S473E or S489D substitution alone suffices to destabilize chromatin anchoring, suggesting both sites converge on the same chromatin-release mechanism, and *in vivo* ChIP-Seq in R1α^WT^ *vs* R1α^cKO^ ovaries revealed significant loss of TRIM28 binding sites in PKA GOF context. Integration of RNA-seq and ChIP-seq data identified 271 DEG in R1α^cKO^ ovaries containing at least one lost TRIM28 binding site, including ovarian and testicular-related genes. BETA analysis showed TRIM28 binding sites lost in R1α^cKO^ were associated with ovarian downregulated genes, supporting TRIM28 as a transcriptional co-activator in 12-week-old-R1α^WT^ ovaries. These observations, together with motif enrichment findings close to TRIM28-bound sites, are consistent with our recent highlights identifying TRIM28 as a central organizer of ovarian TF hubs whose predominant function is to preserve GC identity through stabilization of specific TFs rather than its involvement in H3K9me3-dependent heterochromatin^86^. The demonstration that chronic PKA GOF destabilizes FOXL2–TRIM28 physical interaction in GC— worsened by 12 weeks— provides the functional link between our biochemical findings and the established ovarian identity-maintenance paradigm centered on FOXL2^4^ and its partnership with TRIM28^5^. Recent studies highlight FOXL2 interacts with different binding partners to regulate GC differentiation dynamics ^67^. Given FOXL2 drives supporting gonadal cell differentiation in early ovarian development^69^ acting both as transcriptional activator and repressor^87^, these studies emphasize FOXL2 dual role as differentiation driver and identity-maintenance factor. PLA data showing transient TRIM28–FOXL2 interaction destabilization during gonadotrophin kinetics in WT ovaries extends this to physiological context, suggesting cyclical PKA activation creates transient windows for FOXL2–TRIM28 complex weakening that are reversible under normal conditions but become irreversible under chronic PKA GOF. TRIM28 and FOXL2 cistrome integration identified co-bound genes from which FOXL2 was displaced in R1α^cKO^ ovaries, with 195 genes showing strong transcriptional deregulation. This co-regulated target list, including genes governing follicle plasticity and GC differentiation as well as Sertoli cell signatures, represents the core PKA–TRIM28–FOXL2 axis genomic output. 57% of these shared targets are upregulated and 33% downregulated in both R1α^cKO^ and Trim28^cKO^ ovaries, confirming TRIM28 and FOXL2 co-regulate these genes in the same direction.

Exploration of the R1α-T28^dKO^ model (PKA-GOF + Trim28-deletion) revealed synergistic, accelerated trans-differentiation with pseudo-tubular structures invading the entire ovary by 5 weeks, when R1α^cKO^ ovaries show only modest molecular changes. This synergy demonstrates PKA-GOF targets ovarian identity-maintenance through both TRIM28-dependent and TRIM28-independent mechanisms. First, we identified marked *Amh* as new downregulated gene in R1α-T28^dKO^ ovaries specifically. Since AMH restrains follicular reserve recruitment^78,88^, AMH loss in R1α-T28^dKO^ would amplify GCs subjected to trans-differentiation pressures. Second, *Bhmt* downregulation in the shared R1α-T28^dKO^/R1α^cKO^/T28c^KO^ cluster is intriguing. *Bhmt* is involved in one-carbon metabolism^89,90^, and PKA-driven perturbation could alter methyl donor availability, creating a self-reinforcing epigenetic deregulation cycle. Third, specific deregulation of 55 epigenetic regulators and 72 TFs in R1α-T28^dKO^ ovaries, including SUMOylation targets, reveals chronic PKA activity extends beyond TRIM28 to encompass broad epigenetic and transcriptional regulatory networks.

Together, our data support an integrative model where cyclical gonadotrophin pulses driving folliculogenesis create transient, reversible windows of PKA-mediated TRIM28 phosphorylation, hypo-SUMOylation, and FOXL2–TRIM28 complex destabilization. Under physiological conditions, these transient windows are integral to functional plasticity required for follicle maturation and ovulation. However, when PKA activity becomes chronic—as in R1α^cKO^ model or CNC patients—the transient windows become persistent, the TRIM28– FOXL2 chromatin complex is progressively dismantled, and accumulated SUMOylation-dependent epigenetic identity-maintenance loss culminates in trans-differentiation toward a testicular somatic cell program. This positions SUMOylation dynamics as a central integrator of hormonal signaling and epigenetic identity-maintenance in the ovary, extending the concept of SUMOylation as a guardian of cell identity^11–15,17^ to the reproductive context^82,91^.

## METHODS

### Mice models

*Sf1:Cre*^58,92^, *Prkar1a^fl/f^ ^l^*^58,93^, *Trim28^fl/fl^* ^5^, *Prkaca^+/–^*^58,94^ and *AR^fl/fl^* ^95^ mice have been described previously. Mouse models were enriched with the Rosa26R-mtmg system to follow recombination events. Control animals were *Prkar1a^fl/fl^*littermates, depending on experiments. Mice bred in-house and maintained on a mixed C57Bl/6 genetic background were housed on a 12-h light/12-h dark cycle (lights on at 7:00 am). Mice were fed normal, commercial rodent chow and provided with water ad libitum in a mice facility (approval number: E6301419). After weaning, mice were kept in siblings with a maximum of five animals per cage.

### Samples from patients

Histological slides of ovaries lesions were obtained from 3 CNC patients from the Professor C. Stratakis Service (Section on Endocrinology and Genetics, Eunice Kennedy Shriver National Institute of Child Health and Human Development (NICHD), NIH, Bethesda, Maryland, USA). Human adult healthy testis and ovary sections were used as controls for histological analysis and RNA levels analysis (UM-HuFPT151, CliniSciences). The clinical data summary of CNC patients is provided in Supplementary Table S1. **eCG/hCG treatement.** 100μg of Cetrorelix acetate (C5249, Sigma) were injected intraperitoneally once on female WT OF1 mice, at 22-23 dpn. In addition to Cetrorelix acetate, ovarian stimulation was performed in 23- to 25-day old female mice *via* intraperitoneal injection of 7 IU of eCG/PMSG (equine chorionic gonadotrophin, CHRONO GESTⓇ PMSG 600, MSD) to stimulate follicle maturation followed primed with 7 IU of hCG (human chorionic gonadotrophin, ChorulonⓇ 1500, MSD) 50h later to induce ovulation. For mini-puberty experiments, 100μg of Cetrorelix acetate (C5249, Sigma) or saline buffer (PBS) were injected intraperitoneally twice on prepubertal female mice, at 12 and 13 dpn.

### Hormonal measurements

At the end of experimental procedures mice were killed by decapitation around 8:30 am and trunk blood was collected in vacuum blood collection tubes (VF-053STK, Terumo). Plasma testosterone concentrations were determined using a commercial ELISA kit (LDN ELISA kit AR E-8000). Plasma LH and FSH concentrations were determined using a multiplex assay (MPTMAG-49K, Merck Millipore) as previously described^58^. Further description of methods is provided in Supplemental Methods.

### Cell culture and plasmids

The primary cultures were obtained from ovaries of 3-week-old OF1 mice following the protocol described in Lenie et al. 2004^96^. Briefly, the collected ovaries were punched with a needle 30G to release the granulosa cells, which were then seeds at a concentration of 100,000 cells/ml into 12-well culture plates. The cells were cultured in a DMEM-F12 medium (Thermo Fisher Scientific, Waltham, MA, USA) at 37°C in the presence of 5% CO2, L-glutamine (3 mM), Sodium pyruvate (0,25mM), penicillin /streptomycin (2X) and 1% fetal calf serum. For PKA induction experiments, cells were deprivated of serum one night prior to the addition of forskolin and IBMX (3-isobutyl-1-méthylxanthine) at times and concentrations indicated in the legends of the figures. The identification of primary granulosa cells was confirmed by testing the expression of the *Cyp19a1* and *Star* genes 2 hours after treatment.

For co-transfection experiments, HEK293 cells were cultured on 10 cm cell culture dishes (Millipore Sigma, Burlington, MA) in a DMEM-F12 medium (Thermo Fisher Scientific, Waltham, MA) at 37 °C in the presence of 5% CO2, glycin (1%), penicillin/streptomycin (1%), and 10% foetal calf serum. Cells were seeded at a density of 2×10⁶ cells per 6 cm Petri dish and cultured for 7 hours, after which they were co-transfected using jetPEI (a polycationic transfection agent, Polyplus) and a constant amount of 1.25 μg of the plasmid mixture directing the expression of the human genes of interest (cDNA) under the control of the same CMV enhancer/strong promoter. The parental empty plasmid was added to the mixture to bring the total to 5 μg. All cell harvests and protein extractions were performed 36 hours after transfection. The pCX_Flag_mTRIM28, pcDNA3-TRIM28^S473E^, pcDNA3-TRIM28^S489D^, pcDNA3^S473E-S489D^ and pcDNA3-HA-hSUMO2 expression plasmid were obtained from Dr. Francis Poulat^5,97^. The pCMV-FLAG-SENP2 wt and pCMV-FLAG-SENP2m R577L, K578M (Senp2 LOF) expression plasmid were obtained from Addgene (#18047 and #18713, Dr. Edward Yeh)^98^. The pcDNA3_hPRKACA were obtained from Dr. O’Brien^9^. HEK293T cells were obtained from ATCC (ref CRL-1573).

### Samples preparation prior to LC-MS/MS analysis

HEK293T cells were cultured under standard conditions and subjected to three experimental conditions: (i) untreated control, (ii) stimulation with 50 *μ*M forskolin (FSK) for 30 min, or (iii) transient transfection with a pcDNA-PKAc expression vector for 24 h using Lipofectamine 2000 (Thermo Fisher Scientific). Endogenous TRIM28 was isolated by immunoprecipitation (IP) for 3 hours at 4°C using DynaGreen Protein A/G magnetic beads (Thermo Fisher Scientific), pre-incubated with 2.5 *μ*g of anti-TRIM28 antibody (cat# PA5-27648, Invitrogen) for 1 hour at 4^∘^C, in the presence of a comprehensive inhibitor cocktail (PIC, PhosSTOP, NaF, b-Glycerophosphate, and SUMO proteases inhibitor N-ethylmaleimide (Millipore Sigma) (3.13 mg/mL)). Forty microliters of 50mM NH4HCO3 buffer containing 1µg of either mass spectrometry-grade Lys-C or sequencing-grade Chymotrypsin were added to each sample. Samples were incubated overnight at 37°C for Lys-C or 25°C for Chymotrypsin with shaking at 600rpm. The next day, samples were placed on a magnetic rack, and the supernatant was transferred to fresh tube and evaporated to dryness. Samples were then resuspended in 40 µL of H₂O containing 0.1% FA. For each sample, 10 µL of the resulting solution was loaded onto Evotips and desalted according to the manufacturer’s instructions prior to LC-MS/MS analysis.

### LC-MS/MS acquisition and data analysis

Data acquisition and bioinformatics analysis were performed on the ProteoSeine platform (Université Paris Cité, CNRS, Institut Jacques Monod, F-75013 Paris, France). Samples were analysed using a timsTOF Pro 2 mass spectrometer coupled to an Evosep One system operating with the manufacturer’s 30 samples-per-day (30 SPD) method. Raw MS files were processed using PEAKS Studio version 12 (Bioinformatics Solutions Inc.). Data were searched against the Homo Sapiens SwissProt database (release 2024_01, 20,429 entries). Identifications were filtered using a 1% false discovery rate (FDR) threshold at both peptide and protein group levels. Further description of LC-MS/MS acquisition and data analysis is provided in Supplemental Methods.

### Expression and purification of recombinant TRIM28 proteins

The mouse *Trim28* cDNA encoding wild-type (WT) TRIM28 and the phosphomimetic mutants S473E, S489D, and S473E/S489D was cloned into the bacterial expression vector pET-28a, generating N-terminal T7-His-tagged fusion proteins. Expression constructs were transformed into BL21(DE3) *Escherichia coli* cells (New England Biolabs, Cat. No. C2527H). Further description of methods is provided in Supplemental Methods.

### *In vitro* kinase assay

Prior to kinase assays, equivalent amounts of purified wild-type (WT) and phosphomimetic TRIM28 proteins were verified by SDS–PAGE followed by immunoblotting using an anti-T7 antibody (Bethyl Laboratories, Cat. No. A190-117A), whereas Coomassie Brilliant Blue staining was used to verify protein integrity and purity. Equivalent amounts of recombinant TRIM28 proteins were then quantified again by T7 immunoblotting and imaged using a ChemiDoc Imaging System (Bio-Rad) to enable normalization of kinase activity measurements.

For *in vitro* kinase assays, 1 μg of recombinant TRIM28 protein was incubated in kinase buffer containing 10 mM HEPES (pH 8.0), 10 mM MgCl₂, 10 mM DTT, 1 mg/mL bovine serum albumin (BSA), 50 μM ATP, and 0.1 μCi γ-32P-ATP (3,000 Ci/mmol; Revvity, USA) in the presence of 5 U of recombinant cAMP-dependent protein kinase catalytic subunit (Promega, Cat. No. V516A). Reactions were carried out in a final volume of 50 μL for 10 min at 30°C. Kinase reactions were terminated by the addition of 10 μL of 5X Laemmli sample buffer, followed by heating at 95°C for 5 min. Samples were resolved by SDS–PAGE. Gels were first stained with Coomassie Brilliant Blue to verify equal protein loading and subsequently dried before visualization of radiolabeled proteins using an Amersham Typhoon phosphorimager (Cytiva). ^32^P incorporation was quantified using ImageQuant software and normalized to the amount of recombinant TRIM28 loaded, as determined T7 immunoblotting.

### Chromatin Preparation and Sequential Salt Extraction

HEK293T cells were seeded in 100-mm dishes at a density of 5 x 10^6^ cells per dish and transfected with the indicated plasmids using Lipofectamine 2000 (Invitrogen) according to the manufacturer’s instructions. Cells were harvested 30 hours post-transfection for sequential salt extraction using a modified chromatin fractionation protocol adapted from Porter et al.^99^. All buffers were pre-cooled on ice and supplemented with Protease Inhibitor Cocktail (PIC, Roche) and PhosSTOP phosphatase inhibitors (Roche). Cells were washed twice with 5 mL of ice-cold PBS, and cell pellets were resuspended in 1 mL of Buffer A (25 mM HEPES, pH 7.6, 25 mM KCl, 5 mM MgCl2, 0.05 mM EDTA, 0.1% NP-40, and 10% glycerol). The suspension was rotated for 10 min at 4°C, and nuclei were pelleted by centrifugation at 6000 x g for 5 min at 4°C. A 2X modified RIPA (mRIPA) stock solution (100 mM Tris pH 8.0, 2% NP-40, and 0.5% sodium deoxycholate) was used to prepare 1X working solutions containing stepwise increasing concentrations of NaCl (0, 100, 200, 300, 400, and 500 mM). The nuclear pellet was first resuspended in 200 µL of 1X mRIPA buffer (without NaCl), homogenized by pipetting 15 times, and incubated on ice for 3 min. The mixture was then centrifuged at 6,500xg for 3 min at 4°C to separate the soluble fraction (supernatant) from the residual chromatin pellet. The remaining pellet was sequentially extracted using 200 µL of 1X mRIPA buffers with increasing NaCl concentrations (from 100 mM to 500 mM). At each step, the pellet was resuspended by pipetting 15 times, equilibrated on ice for 3 min, and centrifuged at 6,500xg for 3 min at 4°C to collect the respective supernatant. Following the final 500 mM salt extraction, the remaining insoluble chromatin pellet was solubilized by Benzonase (Sigma-Aldrich) 1 U/µl digestion for 30 min at 30°C to release proteins retained within the nuclease-sensitive chromatin fraction. All collected fractions were mixed with Laemmli sample buffer and boiled at 96°C for 10 min prior to Western blot analysis. PCNA was used as a loading and extraction control, analyzed alongside endogenous TRIM28.

### Histology and proximity ligation assay

Ovaries were fixed in 4% PFA for 24 hours and embedded in paraffin. In all, 5-μm sections were deparaffinised and processed for H&E. For immunofluorescence, deparaffinised slides were submerged in antigen retrieval buffer and boiled for 25min. After being rinsed with 1X PBS and incubated for 30min in H2O2 0,3%, they were blocked for an hour with 2.5% horse serum (Vector) and incubated overnight at 4 °C with primary antibody. After rinsing, they were incubated with ImmPRESS polymer (MP-7401, Vector) for 30 min at room temperature. HRP activity was detected with TSA Alexafluor (Thermo Fisher). Primary antibodies and associated conditions are listed in Supplementary Table S2.

For PLA, blocked slides were incubated overnight at 4°C with indicated antibodies followed by Duolink *in situ* PLA (Sigma-Aldrich) anti-mouse (minus) or anti-goat (minus) and anti-rabbit (plus) probes and detection reagents according to manufacturer’s instructions. Primary antibodies are listed in Supplementary Table S3.

Images were acquired with a Zeiss AxioImager M2 or Zeiss Axioscan Z1 slide scanner and were minimally processed for global levels with ZEN 3.11 blue edition. To quantify the intensity of fluorescence and the number of PLA foci in granulosa cells, at least 6 fields (X20 objective) for the largest sections *per* ovaries and n=3 ovaries *per* condition were imaged using a Zeiss AxioImager M2 microscopy and then analysed with QuPath 0.7.0 software^100^. Statistics were conducted using R language^101^ following the pipeline used in this study^16^. Images settings and processing were identical across genotypes.

### RNA isolation, RTq-PCR and transcriptomic analyses

Total mRNAs were extracted from ovaries using the NucleoSpin RNA kit (Machery-Nagel, Düren, Germany) and reverse transcribed as previously described^92^. Primers used are listed in Supplementary Table S4. Relative gene expression was obtained by the ΔΔCt method with normalization to average expression of *actin*. Ovary gene expression profiles for 4-week-old and 12-week-old *Sf1Cre/+:R1a^fl/fl^*and *R1a^fl/fl^* littermates ; and 5-weeks-old *Sf1Cre/+::R1a^fl/fl^, Sf1Cre/+::Trim28^fl/fl^, Sf1Cre/+::R1a^fl/fl^;Trim28^fl/fl^* and *R1a^fl/fl^ ;Trim28^fl/fl^* were analysed using RNA-sequencing. RNA sequencing, library generation and differential expression genes analysis were performed by the GenomEast platform (IGBMC, Illkirch, France). RNA-Seq libraries were generated according to manufacturer’s instructions from 300 ng of total RNA using the Illumina Stranded mRNA Prep, Ligation kit and IDT for Illumina RNA UD Indexes Ligation (Illumina, San Diego, USA). Further description of library preparation and sequencing methods is provided in Supplemental Methods.

Reads were mapped onto the mm10 assembly of *Mus Musculus* genome using STAR version 2.7.10b. Gene expression quantification was performed from uniquely aligned reads using htseq-count version 0.13.5 with annotations from Ensembl version 102 and “union” mode. Read counts have been normalised across samples with the median-of-ratios method proposed by Benjamini and Hochberg. Differential expressions have been implemented using the Bioconductor package DESeq2 version 1.34.0. Gene Set Enrichment Analysis was performed using the GSEA software version 4.4.0 and plotted using the replotGSEA function from the Rtoolbox package (https://github.com/PeeperLab/Rtoolbox). Gene ontology analysis was performed using g:Profiler. Data visualisation was carried out using R software (v4.1.0)72, Pheatmap package was used for heatmaps, Vennerable for Euler diagrams and ggplot2 for plots.

### ChIP-seq analysis

Dissected ovaries were snap-frozen and stored at −80°C. Frozen ovaries were crushed in liquid nitrogen using a mini-liquid nitrogencooled mortar (Bel-Art ref H37260-0100). Powdered tissue samples were immediately fixed in PBS/1% formaldehyde at room temperature for 20 min. Chromatin shearing was performed using a Covaris S220 Sonicator (Covaris). Chromatin immunoprecipitation was performed according to the antibody parameters used in this study^5^, using the Low Cell ChIP-Seq kit. DNA sequencing, library generation and analysis were performed by the GenomEast platform (IGBMC, Illkirch, France). Each ChIP-seq (TRIM28/FOXL2) library was prepared from a pool of two independent immunoprecipitations (IP), and each IP was prepared with ovaries of 4 different animals. Paired-end sequencing reads were first trimmed for adapter sequences and low-quality bases using Trim Galore (v0.6.10) with a minimum Phred quality score of 20 and a minimum read length of 20 bp. Trimmed reads were then aligned to the mouse reference genome mm10 using Bowtie2 (v2.5.4) in very-sensitive local mode (--very-sensitive-local). Only properly paired reads were retained (--no-mixed --no-discordant). Aligned reads were filtered using SAMtools (v1.18 or later) to remove unmapped reads and to retain only high-confidence alignments with a mapping quality score ≥ 30. Filtered reads were subsequently sorted and indexed with SAMtools. Genome-wide coverage tracks were generated from the resulting BAM files using deepTools bamCoverage (v3.5.x) with a bin size of 10 bp. Coverage was normalized using counts per million mapped reads (CPM), reads were extended to their inferred fragment length. For the peak calling, FOXL2-enriched regions were identified by peak calling using MACS3 callpeak (v3.0.2) with the merged aligned reads as treatment and corresponding input as control for each condition WT and KO separately. Narrow Peaks were called in BAM format using a q-value cutoff of 0.01, without model building (--nomodel). TRIM28-enriched regions were identified using MACS3 in broad peak mode (--broad, q-value < 0.01) by comparing merged ChIP-seq reads to the corresponding input control for each condition WT and KO separately.

Overlap between WT and KO peak sets was computed using BEDTools intersect (v2.31.1) with default parameters. WT and KO specific peaks were identified using the -v option to retain non-overlapping regions in each condition. Peaks were further annotated relative to genomic features using HOMER annotatePeaks.pl (v4.11). Heatmaps were generated using plotHeatmap (v3.5.6) based on matrices computed with computeMatrix (v3.5.6) in reference-point mode centered on Peaks, with 1.5 kb upstream and downstream flanking regions. Signal was binned at 10 bp resolution (--binSize 10), and missing values were treated as zero (--missingDataAsZero). For the visualisation of ChIP-seq peaks, the Integrative Genomics Viewer (IGV) (v2.19.2) was used^102^. Motif enrichment was assessed using XSTREME tool from MEME Suite 5.5.9 software^103^ with the full peak regions for motif scanning.

### Western blot

Proteins were extracted from snap-frozen ovaries in RIPA buffer : TRIS 25 mM, EDTA 1 mM, MgCl2 5 mM, NP40 1%, glycerol 10%, NaCl 150 mM supplemented extemporaneously with phosphatase inhibitors (1 mM Na3VO4, 0.5 mM NaF), protease inhibitors (Roche, Basel, Switzerland), and SUMO proteases inhibitor N-ethylmaleimide (Millipore Sigma) (3.13 mg/mL). For western blot, 30 μg of total denatured proteins were loaded on 10% SDS-page gel, transferred on nitrocellulose and detected with primary antibodies (Supplementary Table S5). Signals were quantified with ChemiDoc MP Imaging System camera system (Bio-Rad) and Image Lab software (Bio-Rad). Accumulation of phosphorylated or SUMOylated proteins was normalised respectively to the accumulation of the corresponding unmodified protein or GAPDH/ACTIN.

### Statistics and reproducibility

Statistical analyses for next-generation sequencing data are described in RNA-seq and ChIP-seq methods section. The other statistical analyses were performed with GraphPad Prism 11 software. Details of individual tests are outlined within each figure legend, including a number of replications performed (n) and the reported error as the Standard deviation (SD). Briefly, normality of populations distribution was assessed with Shapiro & Wilk test for n∈[7,5000] or otherwise Kolmogorov–Smirnov normality test. If data followed a normal distribution, homoscedasticity was estimated with a Barlett test. To compare two populations, unpaired, two-tailed t test was used for normally distributed data with the same variance, Mann–Whitney for non-normal distributions and Welch t test for normally distributed data but with different variances. To compare three or more distributions: one-way ANOVA for normally distributed samples with pairwise multiple t tests or Kruskal–Wallis for non-normally distributed samples with planned comparisons using Dunn’s test to determine the genotype effect or the treatment effect. Crosses on the violin plots represent the mean and lines represent the median. Error bars in barplot represent the SD unless otherwise stated. Immunostaining pictures are representative of a group of at least three replicates.

---

**Supplementary figure S1:**
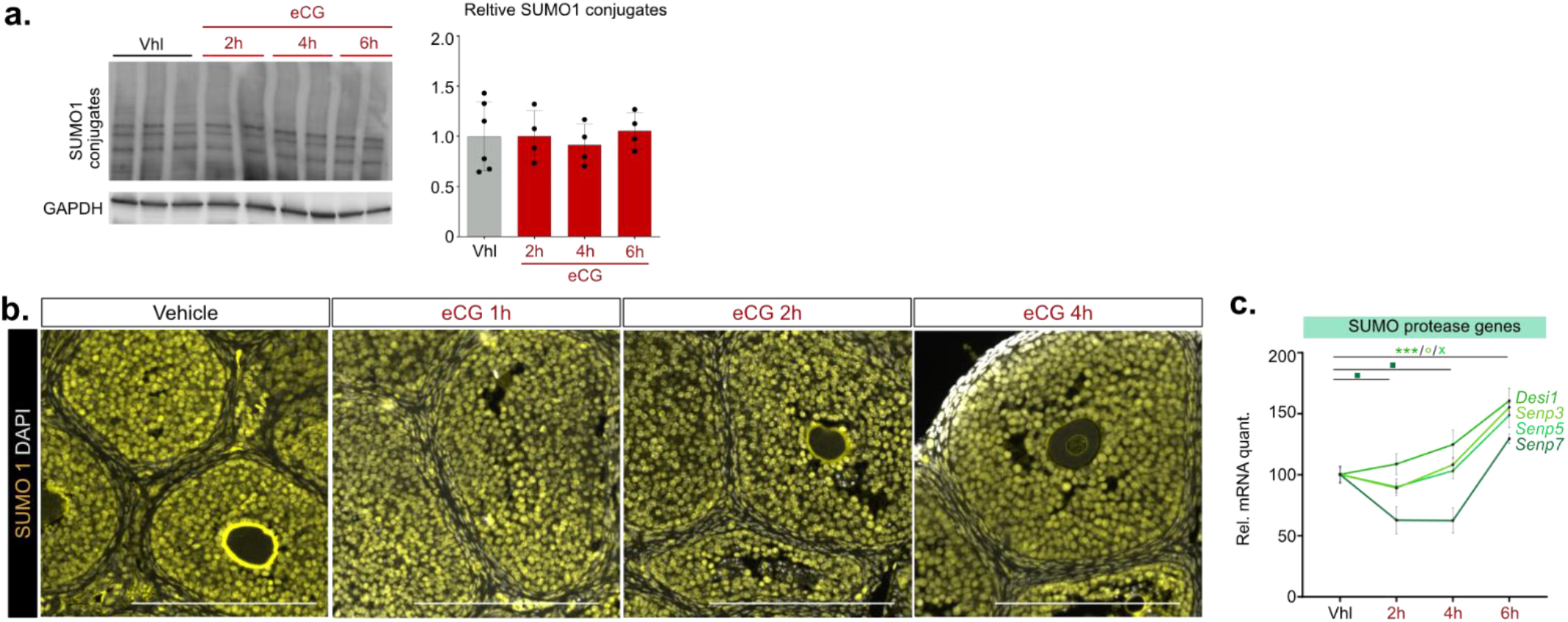
related to figure 1. **a.** Time course effects of gonadotrophins on the accumulation of ovarian SUMO1-conjugated-proteins analysed by Western blot (representative figure) with SUMO1-antibody and SUMO1 signal quantification (histogram). GAPDH signal was used to normalize the quantification of SUMO signals. Statistical analysis was performed using two-sited Mann-Whitney U test. **b.** Immunofluorescence labelling of SUMO1 in WT treated ovaries. Scale bar: 200µm. **c.** RT-qPCR analyses of transcripts encoding the SUMO proteases DESI1, SENP3, SENP5, SENP7 in primary GC. Quantification histograms represent means ± SD. n= 5/6 ovaries *per* genotype. Statistical analyses were conducted by a Kruskal–Wallis test with multiple comparisons using Dunn’s test. \**P*<0.05, \*\**P*<0.01, \*\*\**P*<0.005.

**Supplementary figure S2:**
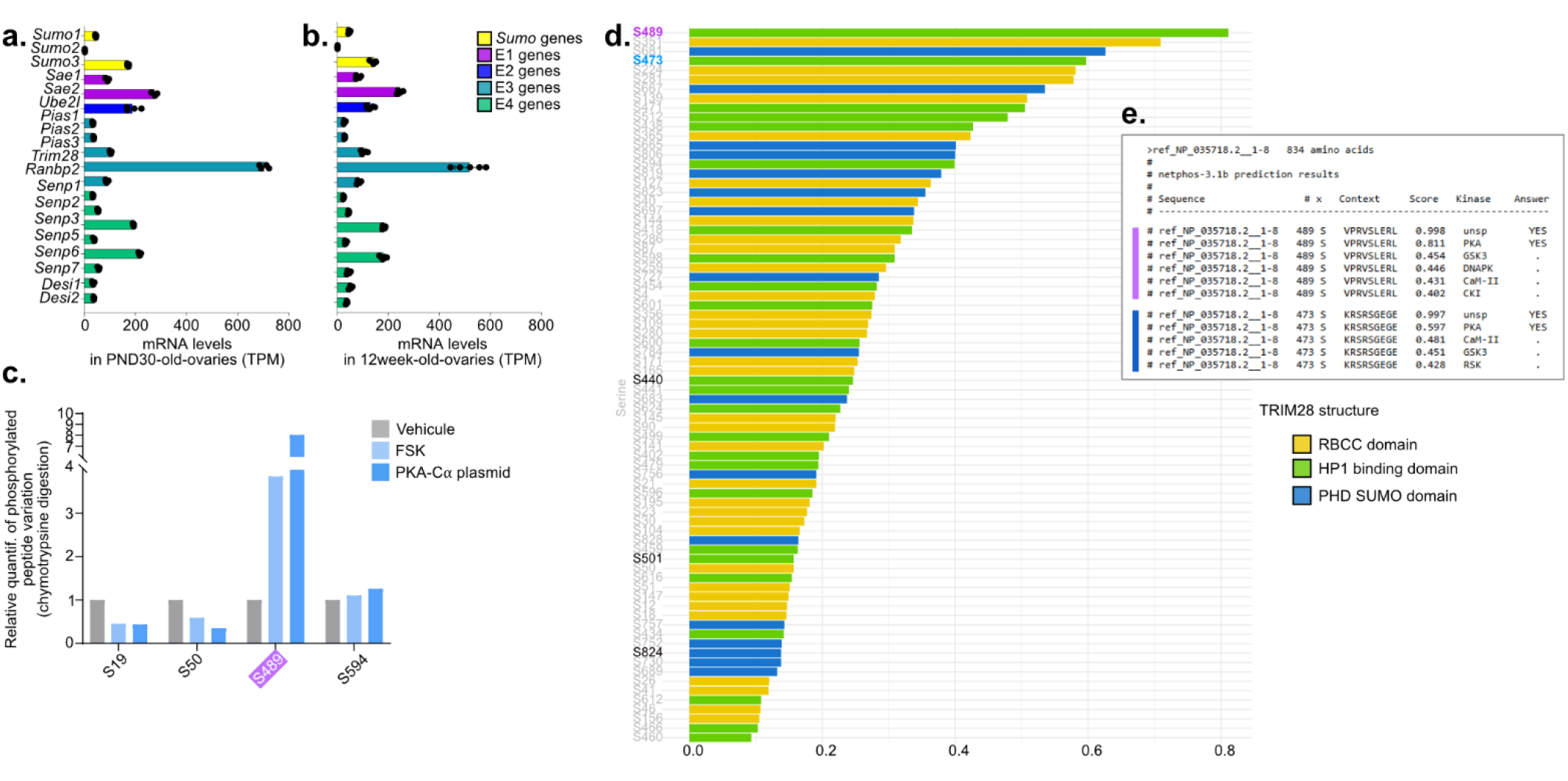
related to figure 2. **a. b.** Relative mRNA abundance of the main actors of SUMOylation (TPM: Transcripts Per Million) extracted from bulk RNA-Seq analyses from ovaries of the PND30 (**a**.) and 12 weeks-old (**b**.) WT ovaries (R1a^WT^ mice) (DPN: day post-natal). **c.** Phosphoproteomic analysis of purified TRIM28. Purified TRIM28 was digested with chemotrypsine and analyzed by LC-MS/MS. Phospho-site assignment was validated using AScore and -10logP confidence scores. Fraction was calculated as phosphorylated peptide intensity/ (phosphorylated + non-phosphorylated peptide intensity). For each condition, a relative quantification of the variation in phosphorylated peptides compared to the vehicle condition was calculated. **d. e.** Prediction score for PKA phosphorylation of the serine residues in TRIM28 from the Netphos analysis. The serine residues phosphorylated by PKA that have been previously experimentally validated by other groups are shown in black^104^.

**Supplementary figure S3:**
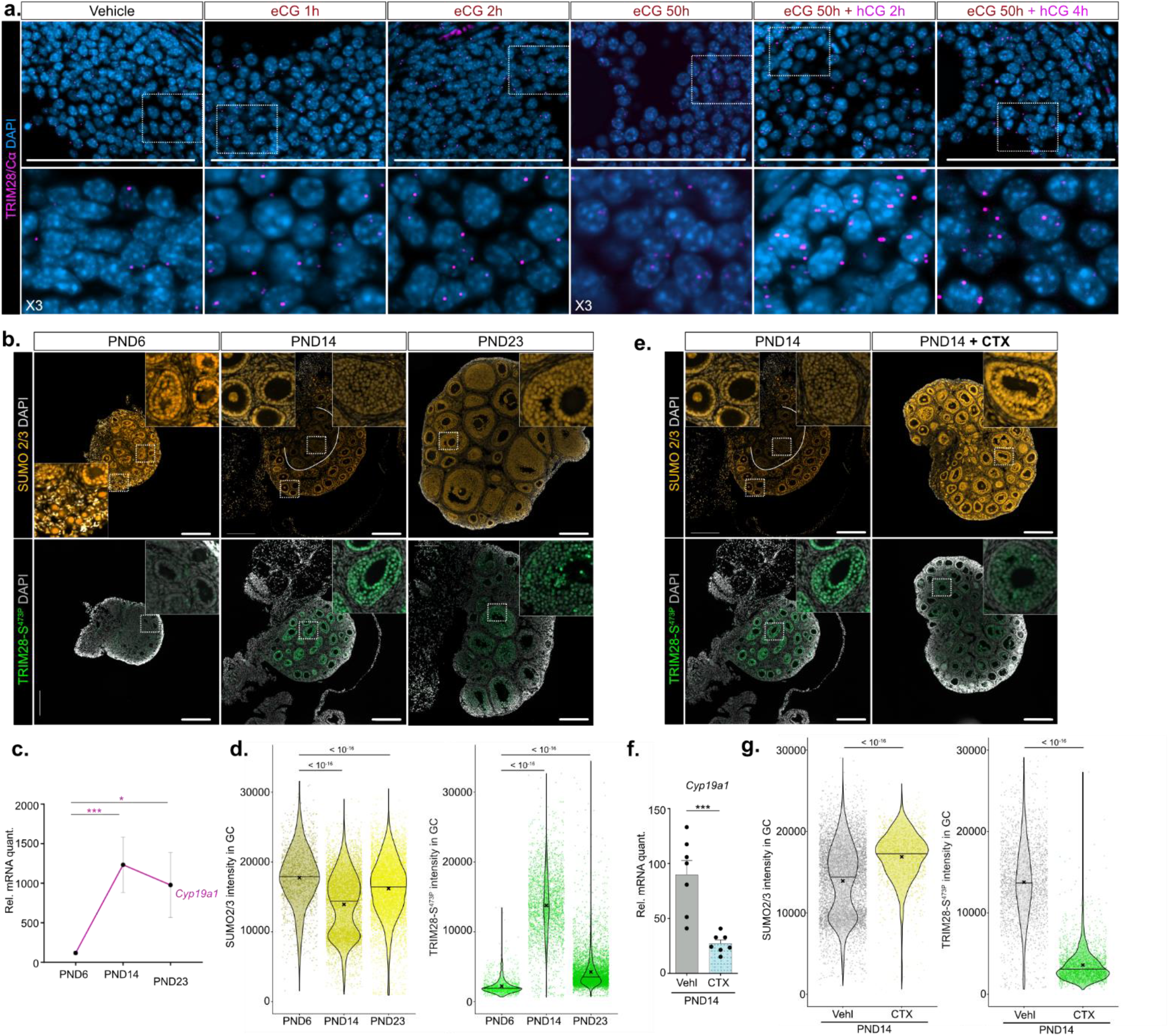
related to figure 3. **a.** Histological images of proximity ligation assay (PLA) of PKA interactions with TRIM28 in the GC nuclei of treated ovaries. As illustrated in inset boxes below, magnification reveals purple dots representing foci of TRIM28-PKA interaction. **b**. Immunofluorescence labelling of SUMO2/3 (yellow) and TRIM28-S^473P^ (green) accumulation in ovaries during the PND6-14-23 stages of the infantile and juvenile periods. Scale bar: 200µm **c.** RT-qPCR analyses of the *Cyp19a1* transcript levels at PND6, PND14, PND23. Quantification histograms represent mean ± SD. n= 7 ovaries *per* stage. Statistical analyses were conducted by a Kruskal–Wallis test with multiple comparisons using Dunn’s test. **d.** Histograms of the quantification of SUMO2/3 and TRIM28-S^473P^ immunostaining signal intensity in GC nuclei from ovaries of 3/4 different mice for the PND 6-14-23 stages of the infantile and juvenile periods. n = 3 ovaries *per* stage. *P* value was determined with a Kruskal–Wallis test with multiple comparisons using Dunn’s test. Scale bar: 200µm. **e.** Immunofluorescence labelling of SUMO2/3 (yellow) and TRIM28-S^473P^ (green) at 14DPN of age in ovaries from mice previously treated with Vehl or CTX at 12DPN and 13DPN. Scale bar: 200µm. **f.** RT-qPCR analyses of the *Cyp19a1* transcript levels in PND14 ovaries from CTX *vs.* Vehl treated mice Quantification histograms represent mean ± SD. n= 7 ovaries *per* treatment. Statistical analyses were conducted by a Kruskal–Wallis test with multiple comparisons using Dunn’s test. **g.** Histograms of quantification of TRIM28-S^473P^ and SUMO2/3 immunostaining signal intensity in GC nuclei from ovaries of 3/4 different mice previously treated either with Vehl or CTX. *P* value was determined with a Kruskal–Wallis test with multiple comparisons using Dunn’s test. \**P*<0.05, \*\**P*<0.01, \*\*\**P*<0.005.

**Supplementary figure S4:**
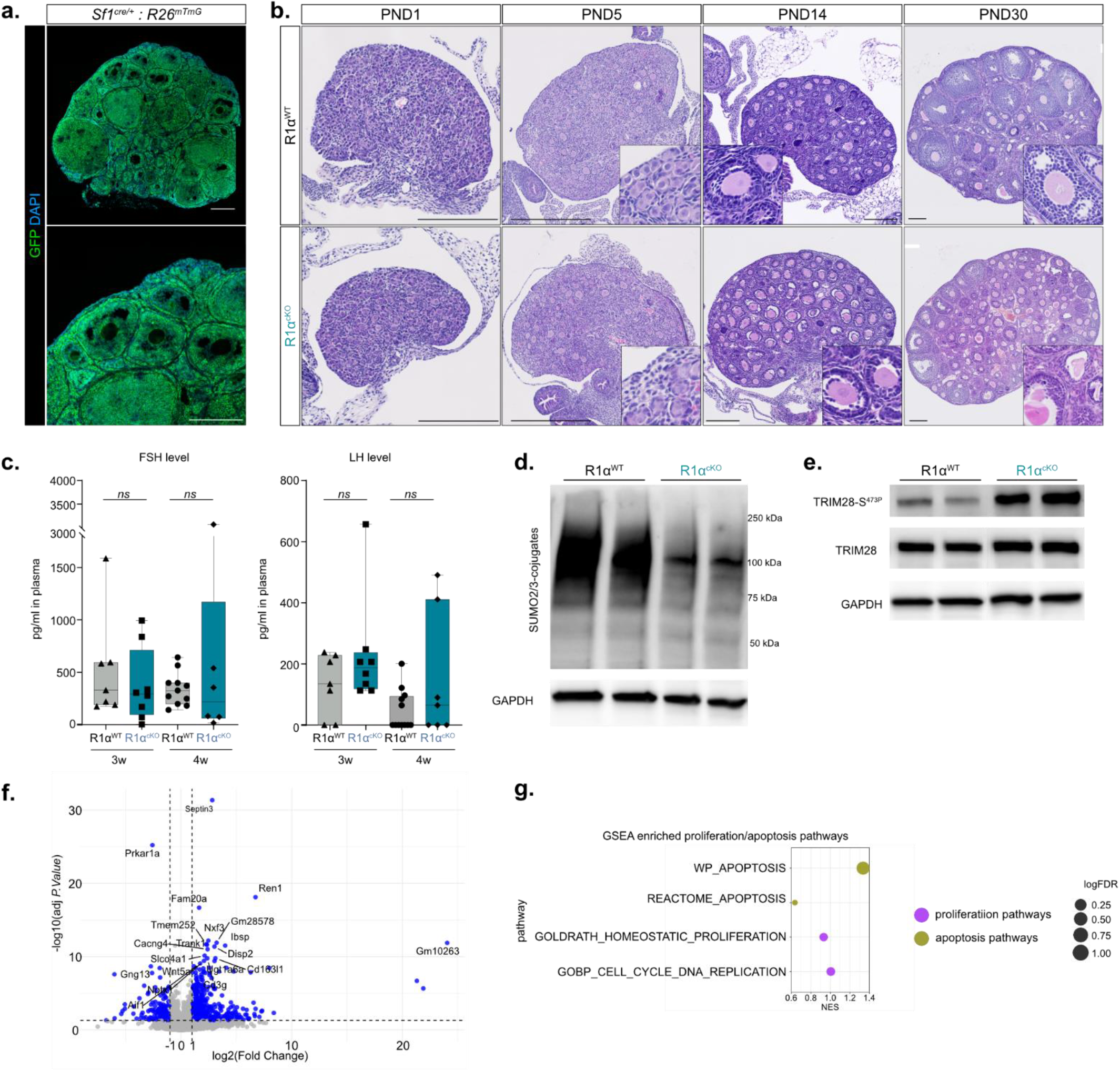
related to figure 4. **a.** Immunofluorescence labelling of GFP (green) in adult *Sf1^Cre/+^ : R26 ^mTmG^* adult ovary attesting wide Cre-mediated recombination activity in ovarian somatic cells . Scale bar: 200µm **b.** Ontogenetic analysis of R1α^cKO^ ovary histology with H&E staining from PND1 to PND30. Scale bar: 200µm. **c.** Plasma FSH and LH concentrations in 3-and 4-weeks-old R1α^cKO^ ovaries compared to R1α^WT^ ovaries. Quantification histograms represent mean ± SD from 6-8 different plasma *per* genotype. Statistical analysis was performed using two-sited Mann-Whitney U test. \**P*<0.05, \*\**P*<0.01, \*\*\**P*<0.005. **d.** Effects of PKA chronic activation on the SUMO2/3-conjugated proteins accumulation from PND30R1α^WT^ and R1α^cKO^ ovaries analysed by Western blot. **e.** Effects of PKA chronic activation on the TRIM28-S^473P^ accumulation from PND30 R1α^WT^ and R1α^cKO^ ovaries analysed by Western blot. **f.** Volcano plot showing the distribution of deregulated genes in PND30 R1α^cKO^ ovaries. **g.** Gene set enrichment analysis (GSEA) of deregulated gene expression data using gene lists associated with proliferation and apoptosis pathways in PND30 R1α^cKO^ ovaries *vs*. PND30 R1α^WT^ ovaries.

**Supplementary figure S5:**
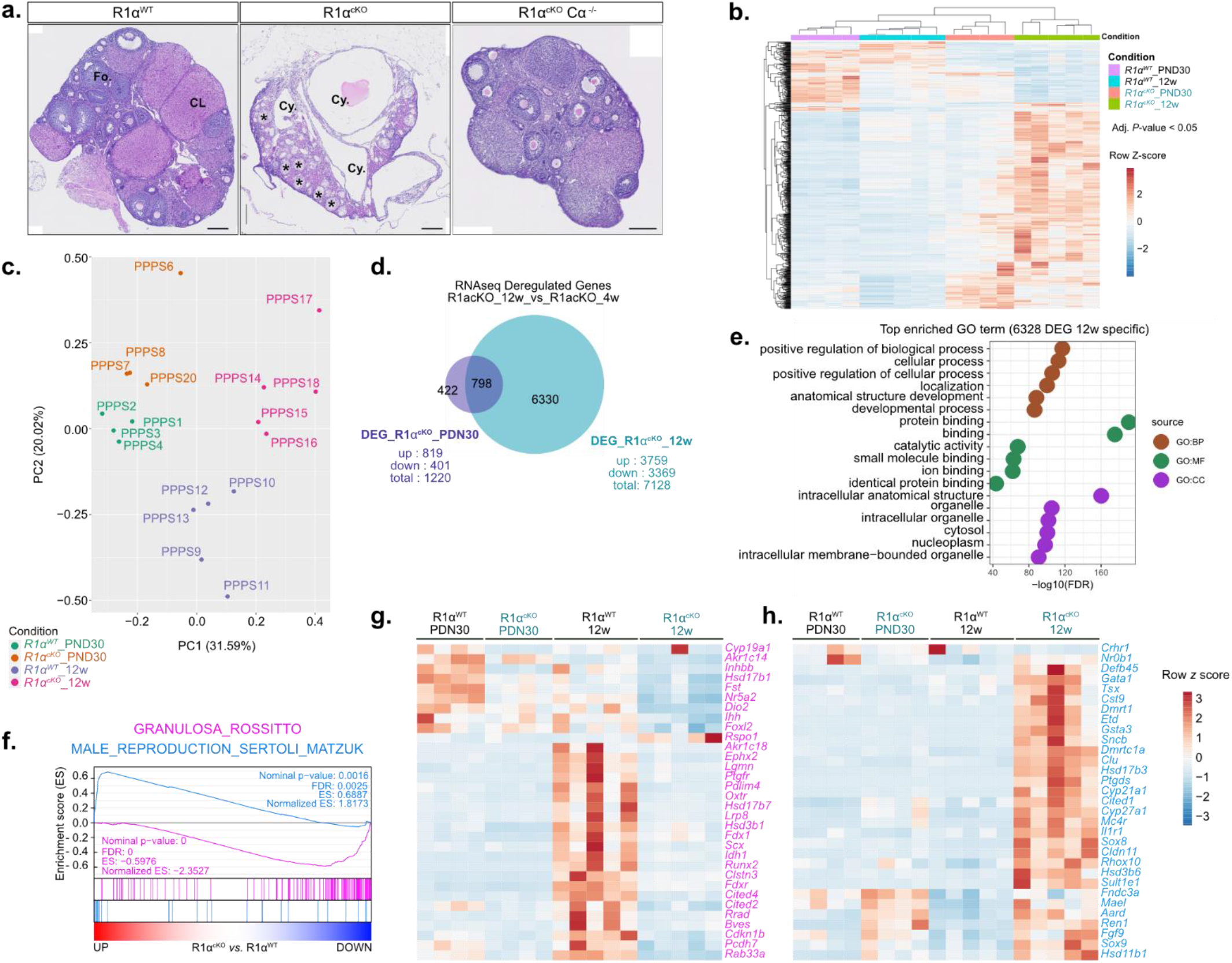
related to figure 5. **a.** H&E staining of R1α^WT^, R1α^cKO^ and R1α^cKO^ Cα ^-/-^12-week-old mice’s ovaries. *\*: pseudo-tubules, CL: corpus luteum, Fo : follicle, Cy : Cysts*. Scale bars: 200 μm **b-c.** Principal component analysis of gene expression in PND30 and 12-week-ol R1α^WT^ and R1α^cKO^ ovaries (**b.**) and associated heatmap (**c.**) representing the median centred expression of dysregulated genes (adj*. P-*value<0.05) determined with Benjamini and Hochberg method, between R1α^WT^ and R1α^cKO^ PND30 and 12-week-old ovaries. **d.** Venn diagram illustrating the commonly deregulated expressed genes (adj*. P-*value<0.05) in both PND30 and 12-weeks-old R1α^cKO^ ovaries **e.** Top GO terms from Gene Ontology functional enrichment analysis based on the common deregulated genes in PND30 and 12-week-old R1α^cKO^. **f.** Gene set enrichment analysis (GSEA) of DEG data using gene lists associated with granulosa/Sertoli-cell-identity markers in 12-weeks-old R1α^cKO^ ovaries compared to the R1α^WT^ ovaries. **g. h.** Heatmap representing the median centred expression of significantly downregulated ovarian expression markers (**g**.) and upregulated testis expression markers (**h**.) in 12-weeks-old R1α^cKO^ ovaries specifically compared to the R1α^WT^.

**Supplementary figure S6:**
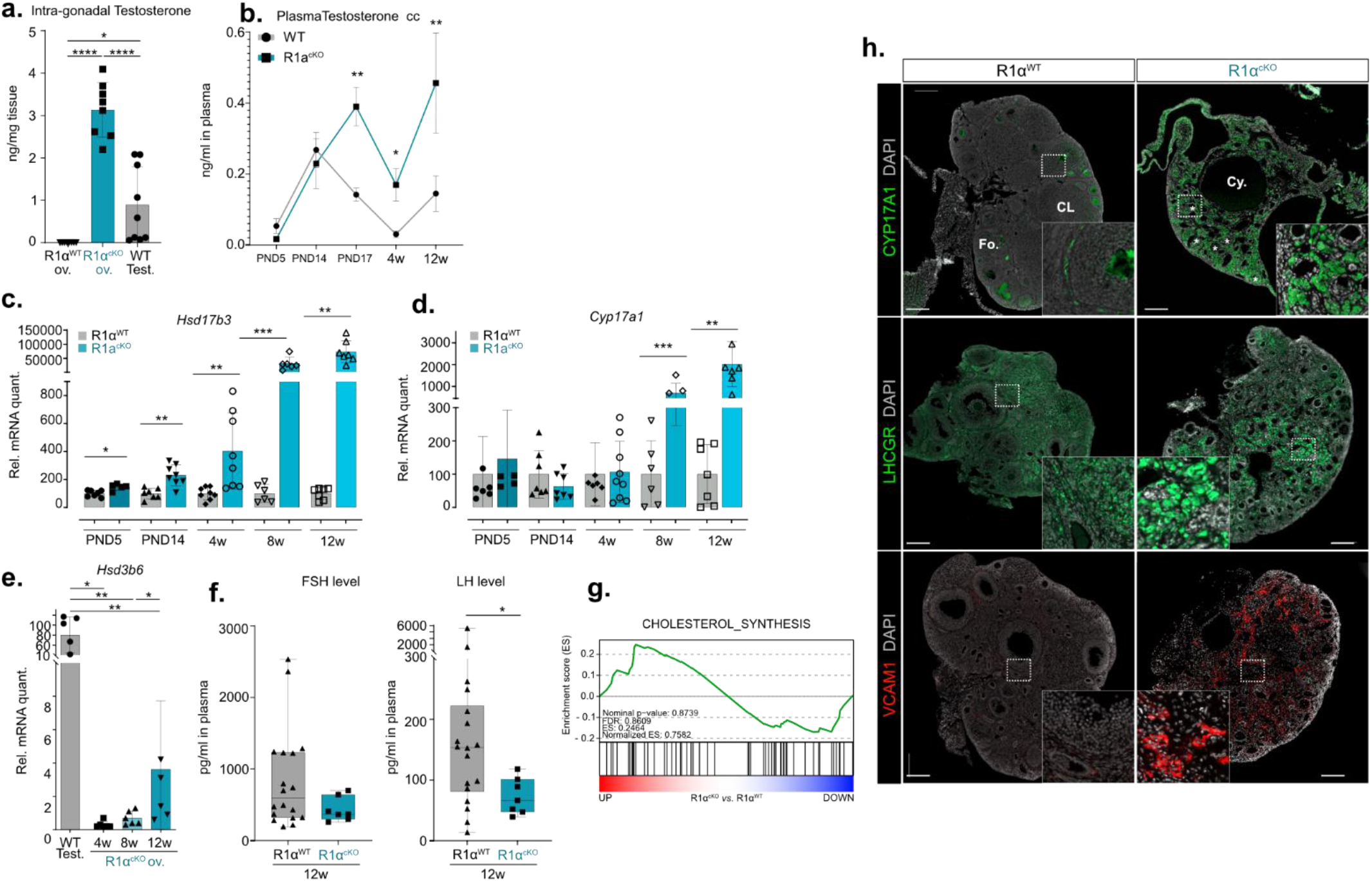
related to figure 5. **a.** Intra-gonadal testosterone concentration in 12-weeks-old R1α^cKO^ ovaries (ov.) compared to 12-weeks-old R1α^WT^ ovaries and testes (test.). Quantification histograms represent means mean ± SD. n= 8/9 mice *per* genotype. *P* values were obtained from one-way ANOVA and adjusted with FDR method. \**P*<0.05, \*\**P*<0.01, \*\*\**P*<0.005. **b.** Ontogenetic analyses of R1α^cKO^ female mice plasma testosterone concentration compared to R1α^WT^ mice from PND5 to 12 weeks of age. Quantification histograms represent mean ± SD. n=5/6 mice *per* genotype. Statistical analysis was performed using two-sited Mann-Whitney U test. \**P*<0.05, ** *P*<0.01, \*\*\**P*<0.005. **c-d.** Ontogenetic RT-qPCR analysis of the *Hsd17b3* (**c.**) and *Cyp17a1* (**d.**) mRNA levels normalized in R1α^cKO^ ovaries compared to R1α^WT^ ovaries from PND5 to 12week of age. Quantification histograms represent means mean ± SD. n=5/6 mice *per* genotype. Statistical analysis was performed using two-sited Mann-Whitney U test. \**P*<0.05, \*\**P*<0.01, \*\*\**P*<0.005. **e.** RT-qPCR analyses of the *Hsd3b6* mRNA levels in PND30, 8- and 12-weeks-old R1α^cKO^ ovaries *vs*. R1α^WT^ testes (test.). Quantification histograms represent mean ± SD. n=5/6 mice *per* genotype. Statistical analysis was performed using two-sited Mann-Whitney U test. \**P*<0.05, \*\**P*<0.01, \*\*\**P*<0.005. **f.** Plasma FSH and LH concentrations in 12-weeks-old R1α^cKO^ mice compared to R1α^WT^ mice. Quantification histograms represent mean ± SD. n=10/12 mice *per* genotype. Statistical analysis was performed using two-sited Mann-Whitney U test. \**P*<0.05, ** *P* <0.01, *** *P* <0.005. **g.** Gene set enrichment analysis (GSEA) of DEG data using gene lists associated with Cholesterol synthesis markers in 12-weeks-old R1α^cKO^ ovaries compared to the R1α^WT^. **h.** Immunofluorescence labelling of CYP17A1 (green), LHCGR (green) and VCAM1 (red) in 12-weeks-old R1α^cKO^ and R1α^WT^ ovaries. *\*: pseudo-tubules, CL: corpus luteum, Fo: follicle, Cy: Cysts.* Scale bars: 200 μm.

**Supplementary figure S7:**
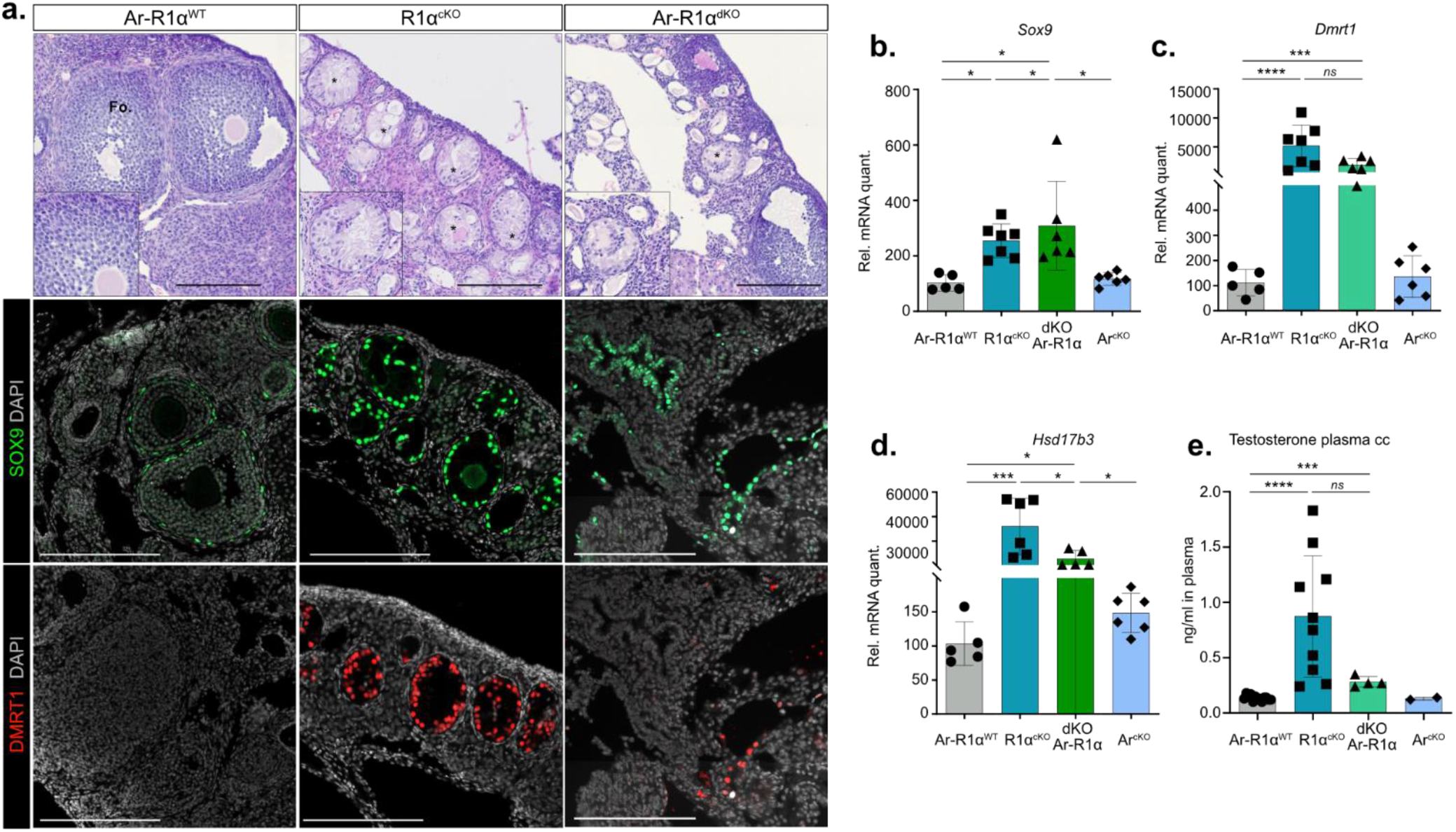
related to figure 5. **a.** H&E staining of R1α^WT^, R1α^cKO^ and Ar-R1α^dKO^ 12-week-old ovaries. Immunofluorescence labelling of SOX9 (green) and DMRT1 (red) in 12-weeks-old R1α^WT^, R1α^cKO^ and Ar-R1α^dKO^ ovaries. *\*: pseudo-tubules, Fo : follicle*. Scale bars: 200 μm. **b. c. d.** RT-qPCR analyses of the *Sox9*, *Dmrt1* and *Hsd17b3* gene expression in 12-weeks-old in R1α^WT^, R1α^cKO,^ Ar-R1α^dKO^ and Ar^cKO^ ovaries. Quantification histograms represent mean ± SD. n=5/7 mice *per* genotype. Statistical analyses were conducted by a Kruskal–Wallis test with multiple comparisons using Dunn’s test. \**P*<0.05, ** *P*<0.01, *** *P*<0.005. **e.** Plasma testosterone concentration in 12-weeks-old in R1α^WT^, R1α^cKO,^ Ar-R1α^dKO^ and Ar^cKO^ female mice. Quantification histograms represent mean ± SD. n=4-10 mice *per* genotype. Statistical analysis was performed using two-sited Mann-Whitney U test. \**P*<0.05, \*\**P*<0.01, \*\*\**P* <0.005.

**Supplementary figure S8:**
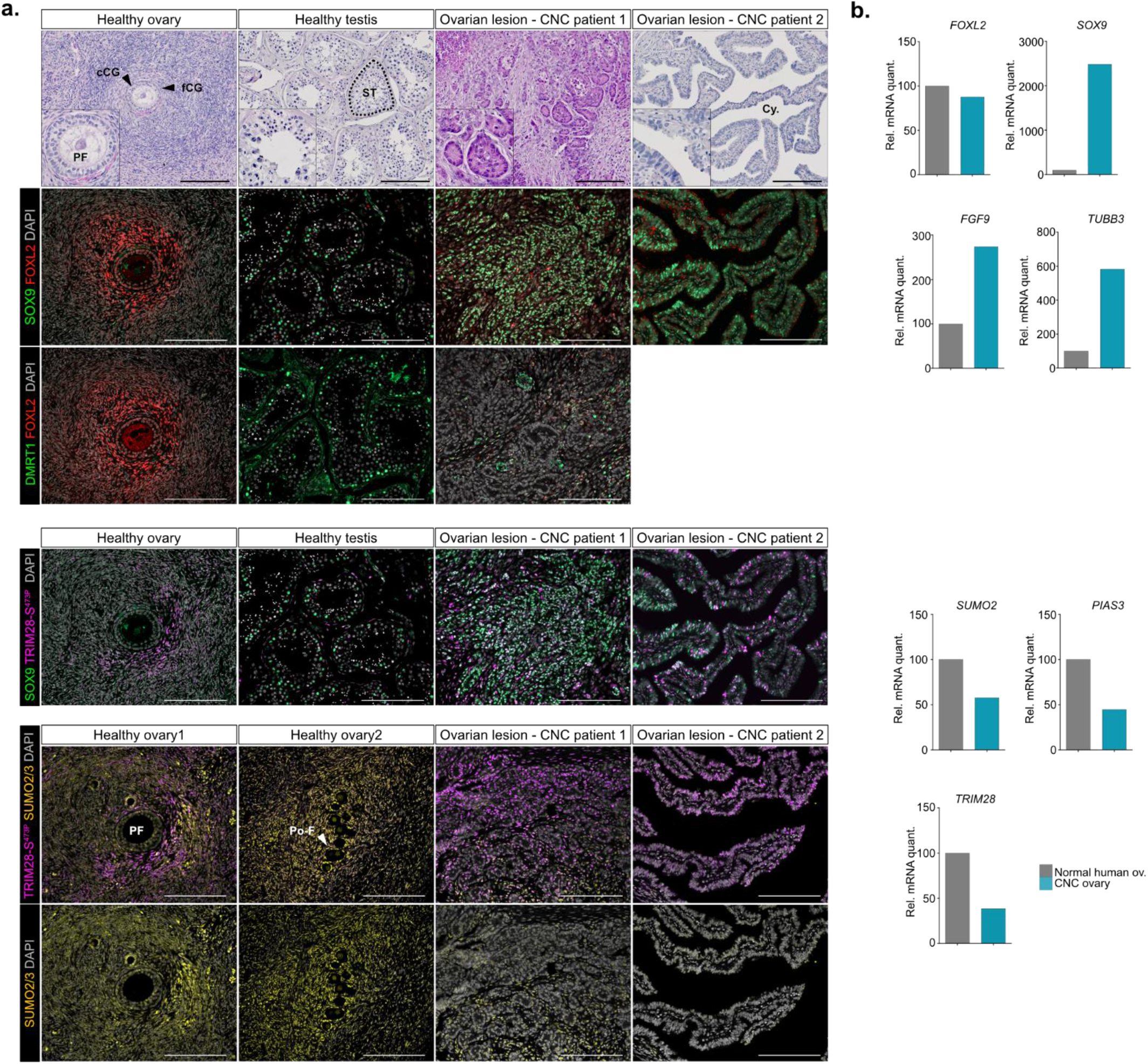
related to figure 5. **a.** Immunofluorescence labelling of SOX9 (green), FOXL2 (red), DMRT1 (red), TRIM28-S^473P^ (purple) and SUMO2/3 (yellow) in ovarian sections of CNC patient 1 and patient 2 (Cy: Cysts), and section of healthy human ovaries sections containing primary follicles (PF, ovary 1) with cubic GC (cGC) and flattened GC (fGC) layers and primordial follicles (Po-F, ovary 2) and testis containing seminiferous tubules (ST). Scale bars: 200 µm. **b.** RT-qPCR analyses of the *FOXL2*, *SOX9, FGF9, TUBB3, SUMO2, PIAS3 and TRIM28* transcripts normalized with respect to *Actb* mRNA levels in CNC patient 3 ovarian biopsy and healthy human ovarian samples.

**Supplementary figure S9:**
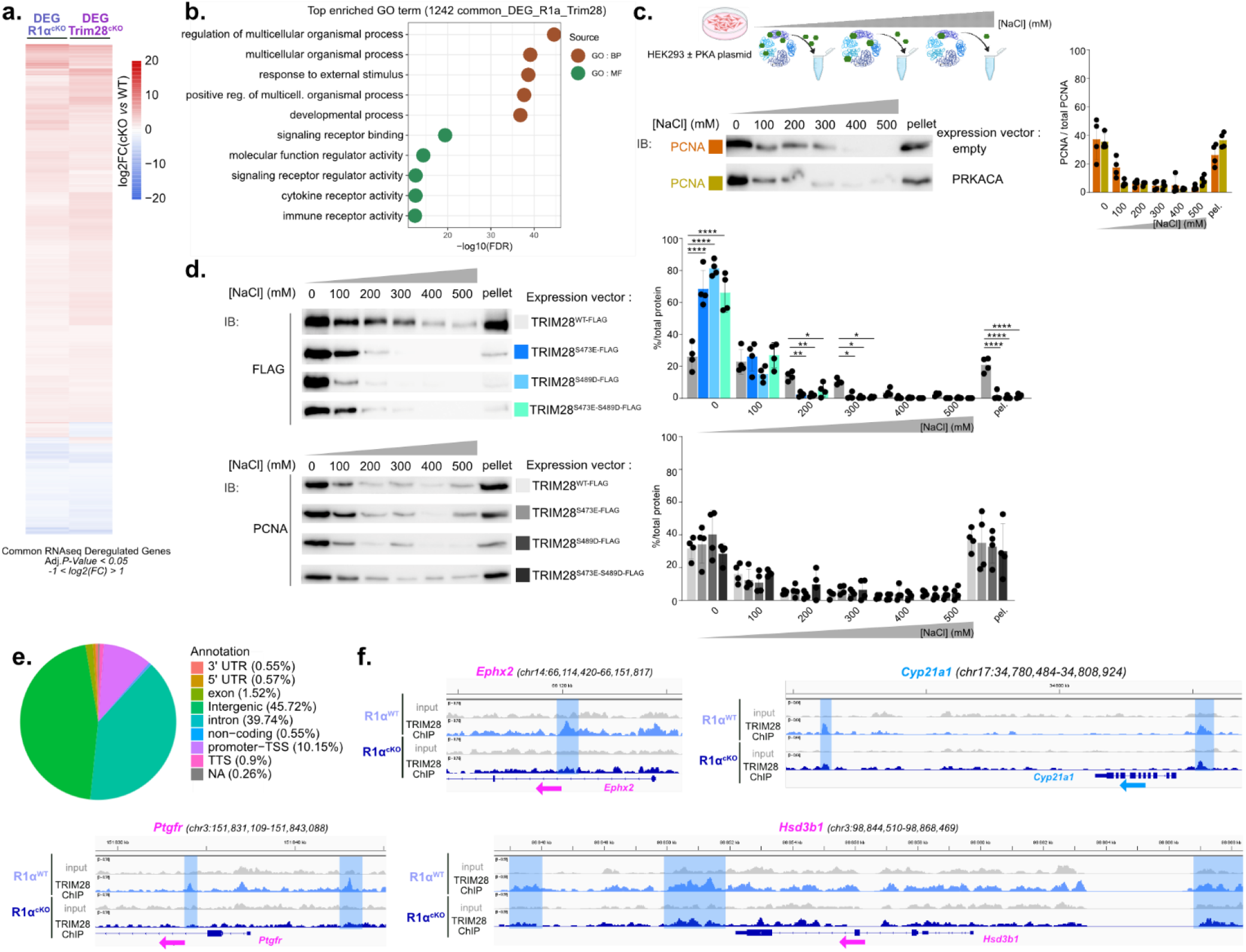
related to figure 6. **a.** Heatmap of the 1242 common deregulated expressed genes (adj*. P-*value<0.05; -1<LOG2FC=1) in R1α^cKO^ and Trim28^cKO^ adult ovaries. **b.** Representative GO biological processes from Gene Ontology enrichment analyses associated to the commonly 1242 deregulated expressed genes. **c.** Measurement of the binding affinity of PCNA to the chromatin extracted from samples mentioned in Fig. 6b. Bottom panel: collected protein fractions analyzed in WB using PCNA antibody. PCNA immunostaining was used as a control for loading and extraction steps. **d.** Measurement of the binding affinity to chromatin of phosphomimetic TRIM28 mutants. Chromatin was isolated as explained in Fig. 6b from transfected HEK293T cells with phosphomimetic TRIM28 mutants (S473E and S489D) and WT-TRIM28 expressing vector. FLAG-tagged TRIM28 WT and the phosphomimetic TRIM28 mutants (S473E and S489D) were analyzed using an anti-FLAG antibody. Phosphomimetic substitutions at Ser473 and Ser489 significantly reduce TRIM28 retention on chromatin. Data are presented as mean ± SEM from n=4 independent experiments. Statistical significance was determined by two-way ANOVA followed by Holm-Šídák’s multiple comparisons test. \**P*<0.05, ** *P*<0.01, *** *P*<0.005, **** *P*<0.0001 **e.** Pie chart mapping in the genome the distribution of the TRIM28 binding sites that were specific to the R1α^WT^ ovaries. **f.** Example of signal intensity tracks showing TRIM28 peaks that were lost in/around genes in R1α^cKO^, and the expression of which was altered in R1α^cKO^ models. Female-specific genes downregulated in R1α^cKO^ ovaries are written in pink in the bottom of the track and male-specific genes upregulated in R1α^cKO^ ovaries are in blue.

**Supplementary figure S10:**
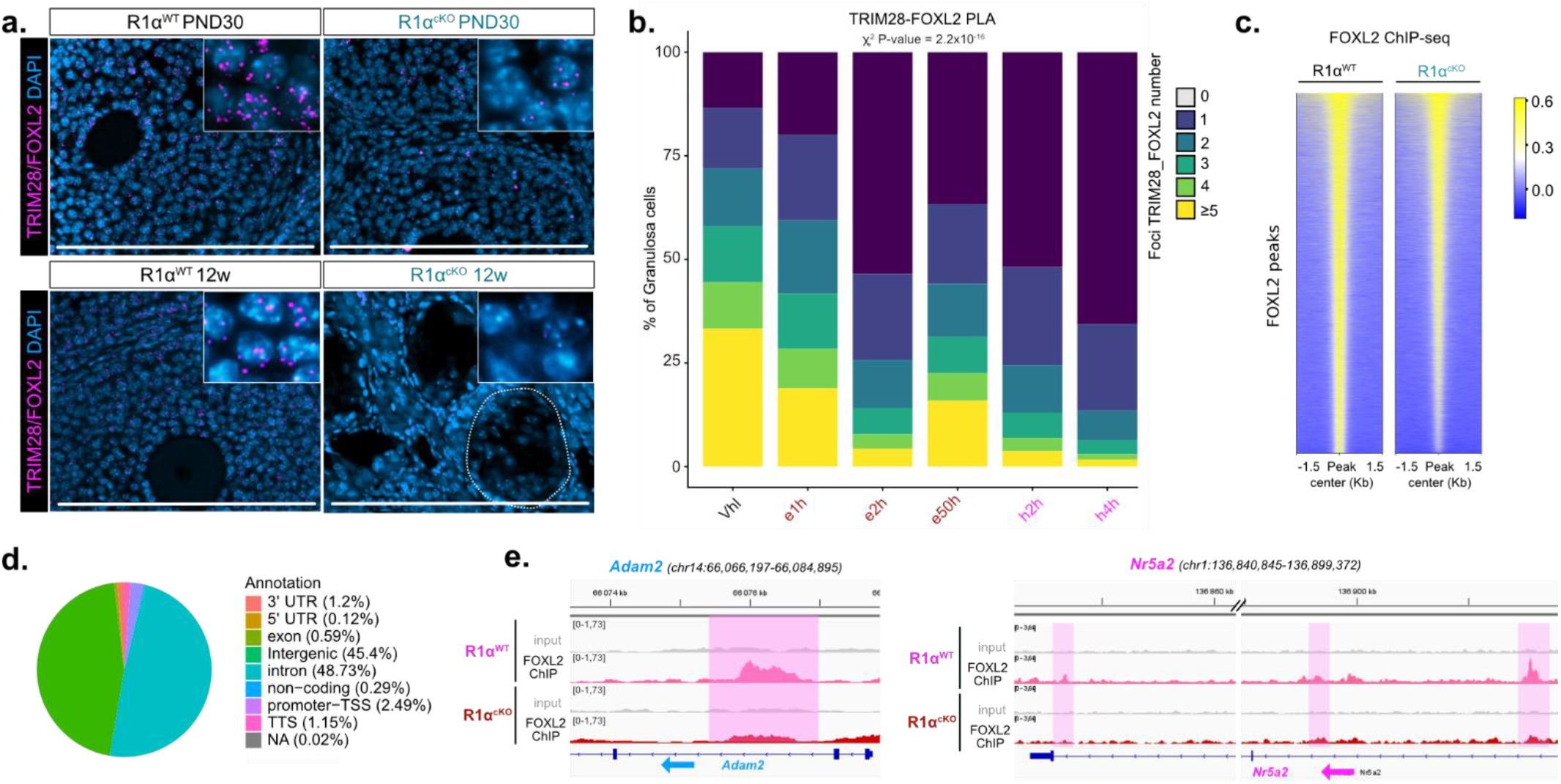
related to figure 7. **a.** Histological images of Proximity ligation assay (PLA) of TRIM28 and its ovarian partner FOXL2 in nuclei of R1α^cKO^ and R1α^WT^ ovaries at PND30 and 12 weeks of age related to Fig. 6a. Scale bar: 200µm. **b.** PLA analysing interactions of TRIM28 with FOXL2 in GC nuclei of gonadotrophin treated WT ovaries. Histograms represent the proportion of GC containing the specified number of dots of interaction *per* nucleus of treated ovaries. To assess whether the distribution of cells differs significantly between the groups a χ² test of independence was assessed. n=3 mice *per* genotype. **c.** Heatmap of normalized FOXL2 ChIP-seq reads around FOXL2 peak center in 12-week-old R1α^WT^ *vs*. R1α^cKO^ ovaries. **d**. Pie chart mapping in the genome the distribution of the FOXL2 binding sites that were specific to the R1α^cKO^ ovaries. **e.** Example of signal intensity tracks showing FOXL2 peaks that were lost in/around genes in R1α^KO^ ovaries and, the expression of which was altered in R1α^cKO^ models. Left panel: male-specific genes downregulated in R1α^cKO^ ovaries. Right panel: female-specific genes upregulated in R1α^cKO^ ovaries.

**Supplementary figure S11:**
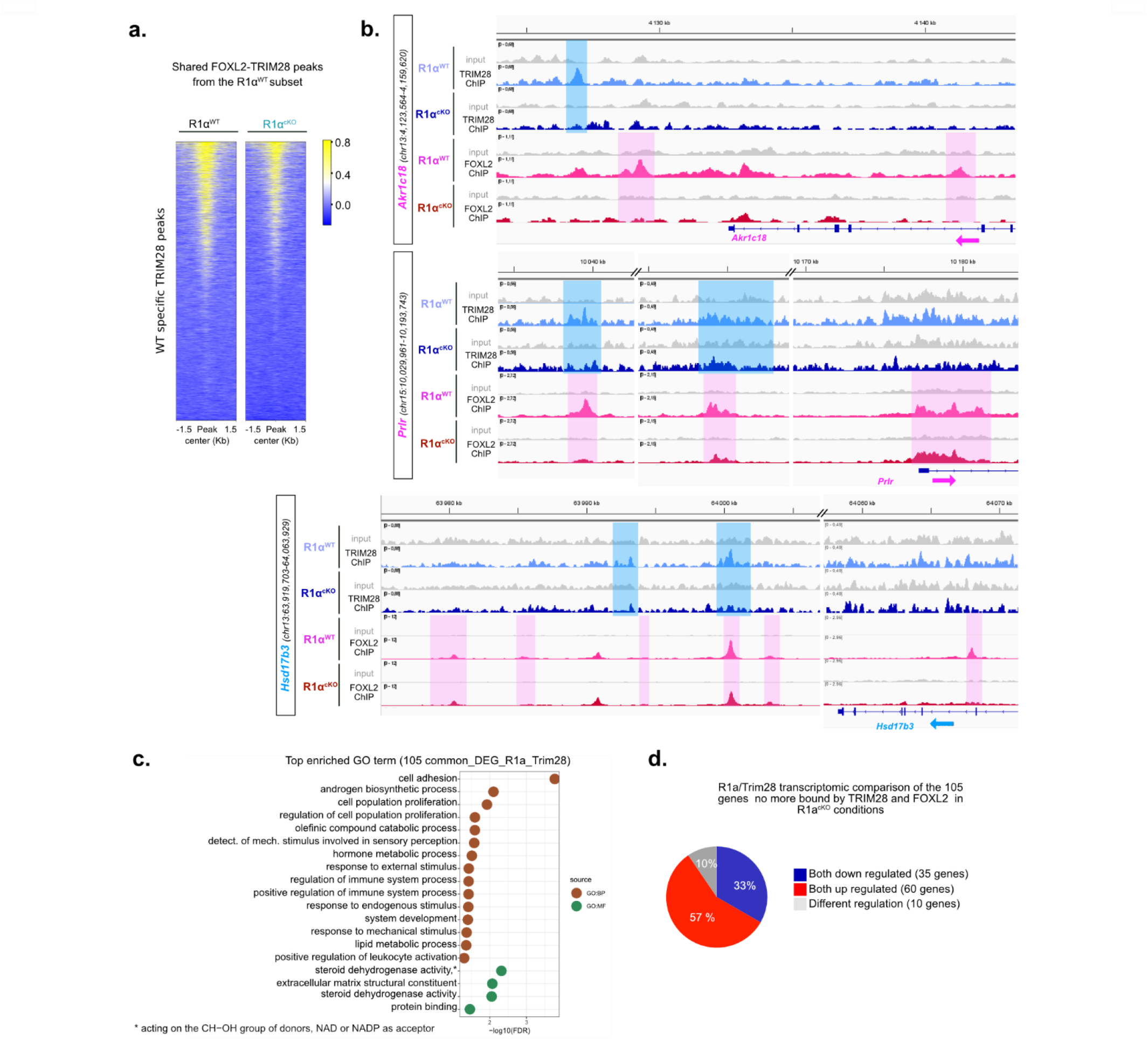
related to figure 7. **a.** FOXL2 genomic localization based on the R1α^WT^ specific TRIM28 peaks in R1α^WT^ and R1α^cKO^ adult ovari s. **b.** Example of signal intensity tracks showing FOXL2 and TRIM28 peaks that are lost in/around genes in R1α^cKO^ ovaries that are deregulated in R1α^cKO^ and Trim28^cKO^. Ovarian specific genes downregulated in R1α^cKO^ are written in pink at the top and bottom of the tracks and male-specific genes upregulated in R1α^cKO^ ovaries are in blue. **c.** Representative GO biological processes from Gene Ontology enrichment analyses associated to the 105 genes whose expression is deregulated in both models (R1α^cKO^/Trim28^cKO^) and which are no more bound by both FOXL2 and TRIM28 in R1α^cKO^ ovaries. **d.** Pie chart showing that most of the 105 genes expression was deregulated in the same way for both models (R1α^cKO^/Trim28^cKO^).

**Supplementary figure S12:**
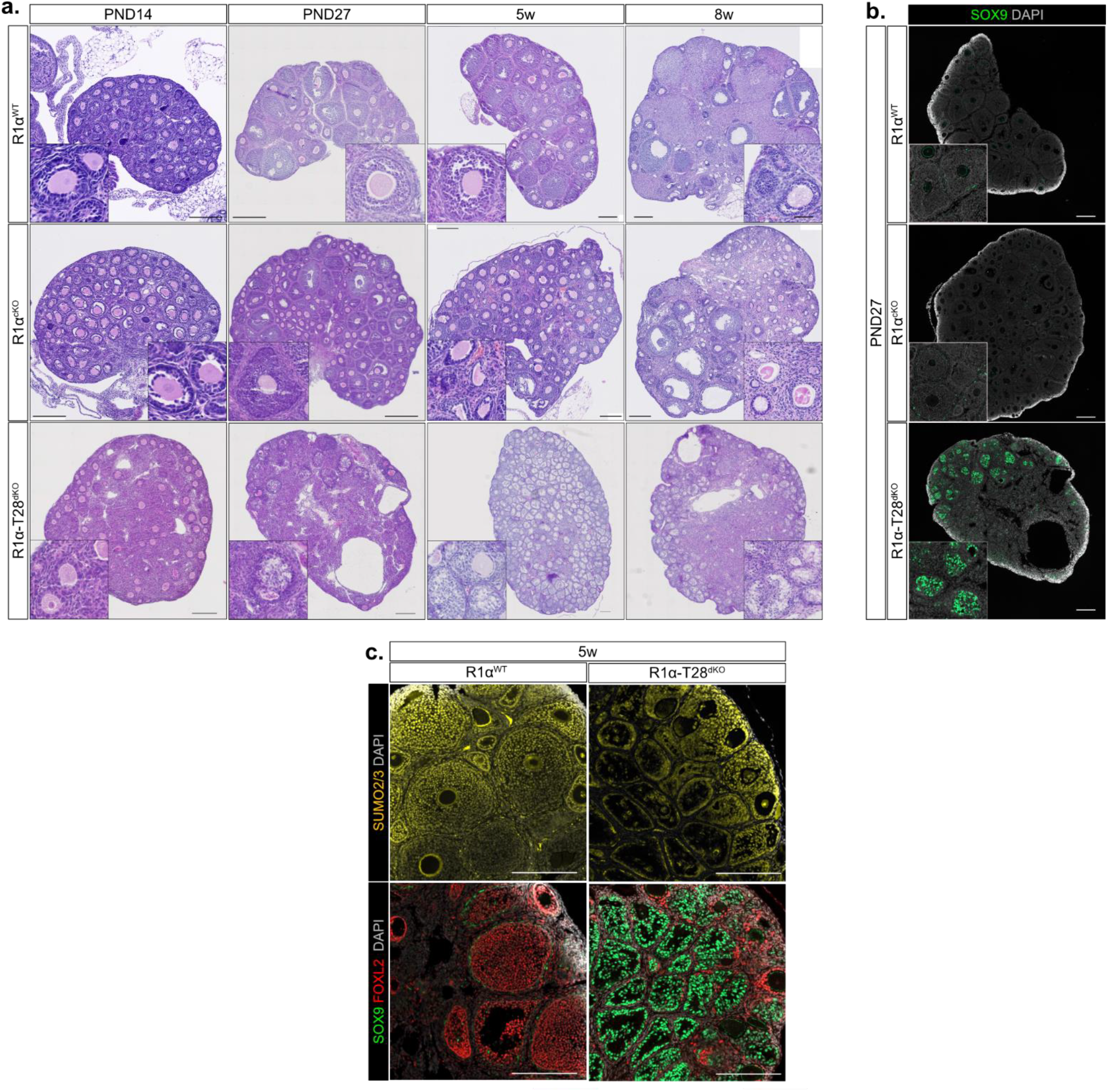
related to figure 8. **a.** Ontogenetic analysis of R1α^cKO^ and R1α-T28^cKO^ compared to R1α^WT^ ^ovary^ histology with H&E staining from PND14 to 8w of age. Scale bar: 200µm. **b.** Immunofluorescence labelling of SOX9 (green) in R1α^WT^, R1α^cKO^ and R1α-T28^dKO^ pubertal ovaries (PND27). Scale bar: 200µm. **c.** Immunofluorescence labelling of SUMO2/3 (yellow), SOX9 (green) and FOXL2 (red) in R1α^WT^ and R1α-T28^dKO^ 5-week-old ovaries. Scale bar: 200µm.

**Supplementary figure S13:**
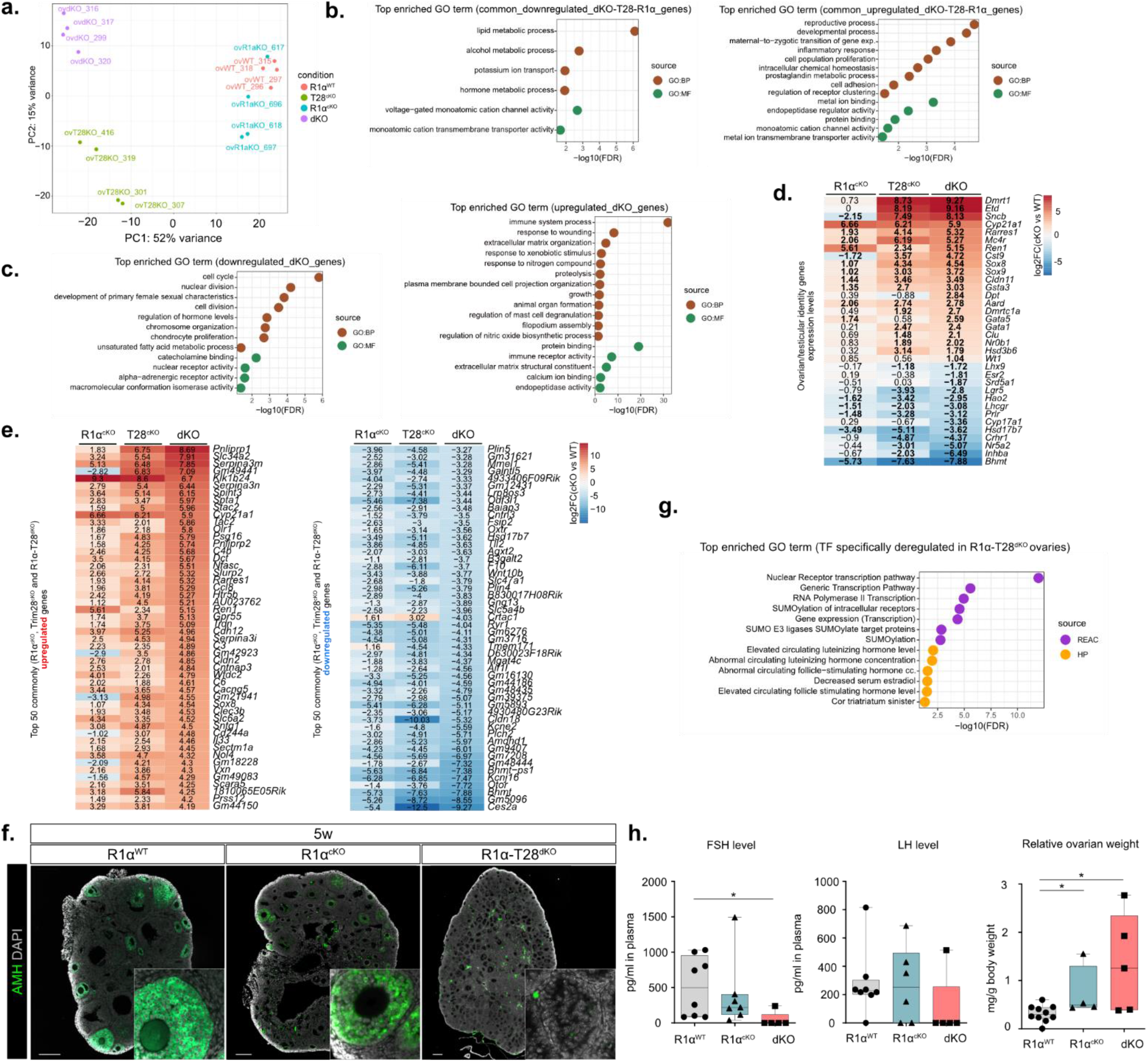
related to figure 8. **a.** Principal component analysis of gene expression in 5-week-ol R1α^WT^, R1α^cKO^, Trim28^cKO^ and R1α-T28^dKO^ ovaries. **b.** Top GO terms from Gene Ontology functional enrichment analysis based on the common downregulated (left panel) and commonly upregulated (right panel) genes in R1α^cKO^, Trim28^cKO^ and R1α-T28^dKO^ 5-week-old ovaries *vs*. R1α^WT^ ovaries. **c.** Top GO terms from Gene Ontology functional enrichment analysis based on the specific downregulated (left panel) or upregulated (right panel) genes in R1α-T28^dKO^ 5-week-old ovaries. **d.** Heatmap representing the log2FC of significantly deregulated genes encoding ovarian/testicular expression markers in 5-week-old R1α-T28^dKO^ ovaries compared to R1α^cKO^ and Trim28^cKO^ ovaries. Genes that are significantly deregulated (adj*. P-value*< 0.05) have their log2FC in bold. **e.** Heatmap of the top 50 most commonly upregulated (left panel) and downregulated (right panel) expressed genes in 5-week-old R1α-T28^dKO^ ovaries compared to R1α^cKO^ and Trim28^cKO^ ovaries (adj*. P-*value<0.05). **f.** Immunofluorescence labelling of AMH accumulation (green) in R1α^WT^, R1α^cKO^ and R1α-T28^dKO^ 5-week-old ovaries. Scale bar: 200µm. **g.** Top GO terms from Reactome and Human Phenotype Ontology databases functional enrichment analysis based on the specific deregulated TFs in R1α-T28^dKO^ 5-week-old ovaries *vs*. R1α^WT^ ovaries (from the list in Fig. 8h). **h.** Left and middle panel: Plasma FSH and LH concentrations in 5-weeks-old R1α-T28^dKO^ female mice compared to R1α^WT^ and R1α^cKO^ female mice. Right panel: Relative ovarian weight (mg *per* g of body weight). Quantification histograms represent mean ± SD. n=5-8 mice *per* genotype. Statistical analysis was performed using two-sited Mann-Whitney U test. \**P*<0.05, \*\**P*<0.01, \*\*\**P*<0.005.

## Acknowledgments

We thank Dr. D. Dufour for his expertise and advice for PLA experiments. We thank Dr .Y. Renaud for management of the GReD institue bioinformatic platform. We thank L. Rigaud, and S. Plantade for animal care. We thank C. Damon-Soubeyrand and S. Bravard (Anipath platform-Clermont) for their technical assistance. We thank staff in the administration and maintenance departments of the GReD Institute.

## Author contribution

F.C., A.M., F.P. and A.-M.L.-M. designed research.

F.P. and F.Ca. provided Trim28-floxed mice.

C.A.S. provided human samples and clinical data.

F.C., L.L.G., I.S.-B., A.-M.L.-M, F.P., S.R., C.D., A.V., P.V., J.-C.P. and L.B. performed experiments.

J.M., S.L.G. and C.K. analysed RNA-Seq and ChIP-Seq data.

F.C., L.L.G., I.S.-B., A.M., A.-M.L.-M. and F. P. analysed data.

A.-M.L.-M and F.C. wrote the paper.

A.M. and F.P. edited the paper.

## Data and materials availability

All data are available in the main text or the supplementary materials. RNA-seq and ChIP-seq data have been deposited in the Gene Expression Omnibus but these data are only accessible to reviewers at the moment.

